# Repetitive Trans-spinal magnetic stimulation Promotes Repair in Inflammatory Spinal Cord Injury Through Sex-Dependent Immune Modulation

**DOI:** 10.1101/2025.10.07.679776

**Authors:** Fannie Semprez, Inès Ziane, Alexandre Du, Inès Istre, Léo Dupuis, Laurine Moncomble, Pauline Neveu, Clémence Raimond, Antoine Fernandes, Jessy Dorange, Quentin Delarue, Nicolas Guérout

## Abstract

Spinal cord injuries (SCI), whether traumatic or inflammatory such as transverse myelitis (TM), are characterized by severe neuroinflammation, demyelination, and long-term disabilities. Current treatments remain limited, highlighting the need for novel non-invasive therapeutic approaches. Repetitive magnetic stimulation (RMS) has emerged as a promising strategy, but its mechanisms and efficacy in inflammatory contexts remain poorly described. Here, we investigated the effects of RMS applied as trans-spinal RMS (rTSMS) in a mouse model of focal spinal cord demyelination induced by lysophosphatidylcholine (LPC). When applied one day after LPC injection, rTSMS reduced inflammation, demyelination, and fibroglial scar formation, while promoting early locomotor recovery in both sexes. In contrast, when treatment was initiated three days after LPC injection, corresponding to the peak of motor deficits, rTSMS conferred tissue protection and functional benefits only in female mice. RNA sequencing analyses revealed sex-dependent immune modulation: in females, rTSMS primarily regulated adaptive T cell–related pathways, whereas in males, it mainly targeted innate immune responses such as neutrophil activity and phagocytosis. Complementary *in vitro* experiments using microglial and macrophage cultures further demonstrated that RMS modulates transcriptomic responses differently depending on cell type and inflammatory state. Specifically, RMS attenuated IL-1–induced pro-inflammatory signaling in macrophages and completely abolished these effects in microglia.

Altogether, our findings establish rTSMS as a non-invasive therapy capable of reducing neuroinflammation and demyelination in inflammatory SCI, with pronounced sex-dependent effects. By uncovering distinct immune pathways engaged in male and female mice, this study provides mechanistic insights into rTSMS action and opens perspectives for its translational use in neuroinflammatory diseases.

## INTRODUCTION

Spinal cord plays a fundamental role in transmitting and receiving motor, sensory and autonomic information from different parts of the body. It is protected by the spine and the meninges, but it can still be damaged. These damages are known as spinal cord injury (SCI), and most often involves physical damage or injury to the spinal cord. SCI causes motor, sensory and autonomic deficits below the site of injury. These are relatively rare conditions, but they remain a major health problem. According to various studies, there are between 250,000 and 500,000 new cases of SCI per year worldwide, with an average incidence in developed countries of 22.5 cases/million population ^1,2^. In 90% of the cases, the injuries are of traumatic origin. As a result, most of these injuries have a common aetiology and are usually caused by preventable events: road accidents, falls, sports activities or violence ^3,4^. As mentioned above, although the majority of SCI are caused by trauma, there are also rare non-traumatic SCI caused by inflammatory diseases or tumours. Inflammatory SCI can be either focal or, conversely, more diffuse. Among focal forms, transverse myelitis (TM) is a rare non-traumatic SCI characterized by inflammatory demyelination of the spinal cord. It disrupts central nervous system–periphery communication, resulting in motor, sensory, and sphincter deficits. Inflammatory processes damage both myelin and axons, affecting immune cells, oligodendrocytes (OLs), and occasionally neurons. TM can occur at any age, with incidence peaks between 10–19 and 30–39 years, and affects males and females equally ^5^. Some cases may result in neuronal loss in severe presentations. In the case of TM, inflammation is highly localised on one level, with thoracic level accounting for 70% of the cases ^5^. This specific localisation of the SCI leads to the same symptoms as a traumatic thoracic SCI. There are many causes of this condition, including infectious and autoimmune diseases ^6^. TM can be also found as a common symptom in a number of CNS inflammatory diseases such as multiple sclerosis (MS), neuromyelitis optica spectrum disorder and myelin-oligodendrocyte glycoprotein antibody-associated disease ^7^. TM causes are most often idiopathic, accounting for 65% of cases, but cases of TM can occur after a viral infection ^5^. The recent Zika and Covid 19 virus pandemics have led to post-infectious TM in some patients ^8,9^.

To date, no specific therapeutic strategy can be proposed to TM patients. The only known treatments are plasma exchange, intravenous steroid injections and immunomodulatory therapies, which only partially improve patients’ daily lives and only for a limited period ^6^. In patients with TM, 33% will have a complete recovery, 33% will have a partial recovery and 33% will have permanent sequelae with no functional recovery ^10^.

Given that certain cellular and molecular mechanisms, such as inflammation and demyelination, are shared between SCI and TM, we hypothesize that treatments previously shown to be effective in the context of SCI may also hold therapeutic potential for TM such as neuromodulation and in particular repetitive magnetic stimulation (RMS). Originally, transcranial RMS is a medical technique which, by generating an electrical current, stimulates or inhibits certain areas of the brain. This technique has already been used clinically to treat neuropsychiatric illnesses, and has a promising future as a treatment for strokes and Parkinson’s disease ^11–13^. Despite the obvious beneficial effects, the associated mechanisms are not fully understood. Our team has focused on focal repetitive trans-spinal magnetic stimulation (rTSMS) (directly and non-invasively at the site of injury) in the context of traumatic SCI in mice in order to understand the underlying molecular and cellular mechanisms ^14^. This previous study shows the ability of rTSMS to modulate spinal cord scar after injury. Motor tests were carried out and showed an improvement in the sensory-motor functions of the mice after treatment with rTSMS. In addition, we were able to demonstrate that this technique reduces demyelination and improves neuronal survival. These results demonstrate the modulating capacity of rTSMS on the various aspects of spinal cord scar in mice ^15^. Furthermore, in a non-lesional context, another team recently demonstrated that rTSMS modulates the proliferation and differentiation of OLs in the mouse brain ^16^. Additionally, it has been shown that RMS can modulate the polarisation of macrophages towards an anti-inflammatory phenotype ^17^.

Based on these findings, we hypothesized that RMS could exert beneficial effects in a focal demyelination model mimicking TM in mice.

## MATERIALS AND METHODS

### Animal experimentation

#### Animal care and use statement

The experimental protocol was designed to minimize pain and discomfort for animals. All experimental procedures adhered to the European Community guidelines on the care and use of animals (86/609/CEE; Official Journal of the European Communities no. L358; 18 December 1986), the French Decree no. 97/748 of 19 October 1987 (Journal Officiel de la République Française; 20 October 1987), and the recommendations of the local ethics committee (#44348).

#### Animals

Mice of both sexes were housed in conventional, secure rodent facilities (two to five animals per cage, sexes separated) under a 12-h light/dark cycle with *ad libitum* access to food and water. A total of 210 adult Wild-type (WT) C57BL/6 mice (8–12 weeks old; mean body mass ≈ 20 g for females, 25 g for males) from Janvier Labs (Le Genest-Saint-Isle, France) were used.

Each experimental group was balanced for sex (50% male / 50% female) and two main study groups were organized as follows (Figure 1A):

1. LPC: animals receiving LPC (lysophosphatidylcholine or lysolecithin) injections in the spinal cord in order to induce non-traumatic SCI.
2. LPC + rTSMS: animals receiving LPC injections in the spinal cord and being treated with rTSMS

**Figure 1:**
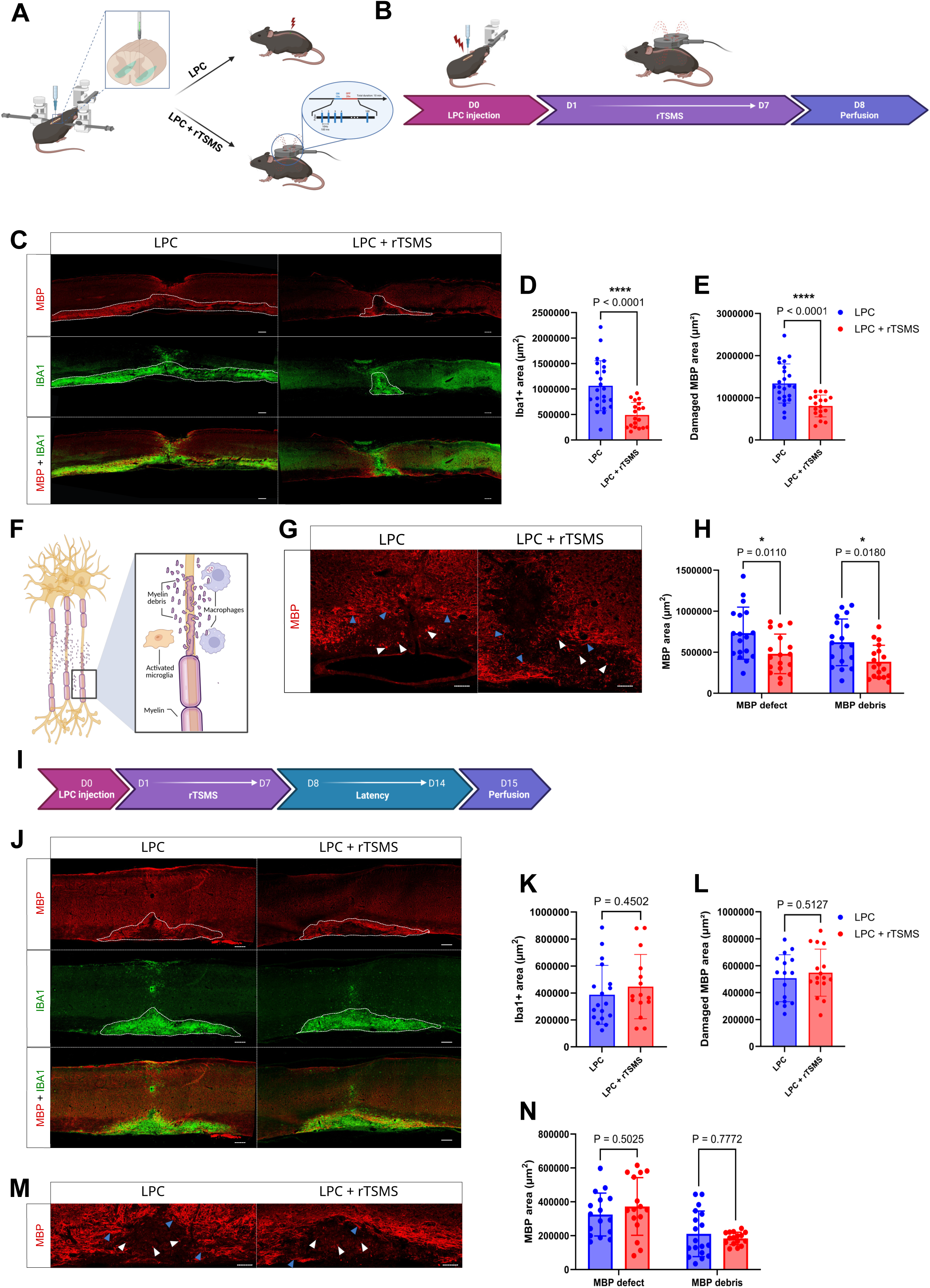
rTSMS reduces demyelination and inflammation in LPC model, when applied at the presymptomatic stage. **A.** Schematic representation of LPC injection surgery and rTSMS treatment in both male and female mice. **B.** Experimental design: on day 0, all mice received an LPC injection; half of them underwent rTSMS treatment from D1 to D7, and all mice were perfused at D8 for histological analyses. **C-H.** Analysis of the effects of rTSMS on inflammation and demyelination processes 8 days after LPC injection with LPC group in blue and LPC + rTSMS group in red. **C.** Representative pictures of sagittal spinal cord sections of LPC and LPC + rTSMS groups 8 days after LPC injection. Sections were stained with anti-MBP (in red) and anti-Iba1 (in green) antibodies for Iba1 and damaged MBP area analyses. The scale bar represents 200 µm **D**. Quantification of Iba1+ area. **E.** Quantification of damaged MBP area (MBP defect + MBP debris areas). **F.** Schematic representation of the different demyelination processes analyzed, including the presence of myelin debris and myelin defects (MBP-area). **G.** Representative pictures of sagittal spinal cord sections of LPC and LPC + rTSMS groups 8 days after LPC injection. Sections were stained with anti-MBP (in red) antibody to analyse MBP defect and MBP debris areas. The white arrows indicate myelin debris, and the blue arrows indicate axons undergoing demyelination. The scale bar represents 100 µm **H**. Quantification of MBP defect (MBP-area) and MBP debris area. **I.** Experimental design: on day 0, all mice received an LPC injection; half of them underwent rTSMS treatment from D1 to D7, and all mice were perfused at D15 for histological analyses. **J-N.** Analysis of the effects of rTSMS on inflammation and demyelination processes 15 days after LPC injection with LPC group in blue and LPC + rTSMS group in red. **J.** Representative pictures of sagittal spinal cord sections of LPC and LPC + rTSMS groups 15 days after LPC injection. Sections were stained with anti-MBP (in red) and anti-Iba1 (in green) antibodies for Iba1 and damaged MBP areas analyses. The scale bar represents 200 µm **K**. Quantification of Iba1+ area. **L.** Quantification of damaged MBP area (MBP defect + MBP debris areas). **M.** Representative pictures of sagittal spinal cord sections of LPC and LPC + rTSMS groups 15 days after LPC injection. Sections were stained with anti-MBP (in red) antibody to analyse MBP defect and MBP debris areas. The white arrows indicate myelin debris, and the blue arrows indicate axons undergoing demyelination. The scale bar represents 100 µm **N**. Quantification of MBP defect (MBP-area) and MBP debris area. N = 17-23 mice in the LPC group and N = 17-19 mice in the LPC + rTSMS group. Statistical analyses were performed using Unpaired t-test (*=P≤ 0.05 and ****=P≤ 0.0001).

#### Surgical procedure and LPC injections

As the precise causes of the onset of TM are still unknown, there is currently no experimental model mimicking the effects of this disease in animals. As a result, the TM development model aims to reproduce the symptoms of demyelination and inflammation of the spinal cord. Focal demyelination models have been studied for many years, and one simple and reproducible model uses LPC ^18,19^. LPC is a detergent that induces destruction of the myelin sheaths, leading both to loss of sensory-motor function in the animals, and to a major inflammatory reaction at the injection site. LPC injection is a good model to mimic TM effects.

All the mice received a premedication of buprenorphine (0.03 mg/mL; 3 µl/g of body weight). The surgical procedure was performed as described by Hérent *al*. ^20^. Briefly, a 2 cm skin incision was performed dorsally, and the exposed spinal column was fixed to a stereotaxic frame using two holders on the left and right sides to minimize movements. Vertebral spinous processes were used as landmarks to target specific segments. Then, two lidocaine injections were administered around the T11 and L1 vertebrae. A small incision of the ligamentum flavum allowed access to the spinal cord. A pulled glass pipette connected to a motorized syringe pump injector (Legato 130, KD Scientific, customized by Phymep, France) was positioned into the ventral funiculus of the L2 (second lumbar segment, between the 11th and 12th vertebral bodies). Demyelination was induced by two bilateral injections of 1µl of 1% LPC (10 mg/ml, Sigma-Aldrich, ref L4129, St. Quentin Fallavier, France) at 0.3 mm lateral to the midline and 1 mm depth, targeting both right and left ventral funiculi, as described by Bardy-Lagarde *et al*. ^21^. Injections were carried out using a 20 μm diameter capillary at a rate of 0.20 μL/min. Before and after each injection, the pipette was maintained in place for 5 minutes and then slowly withdrawn. Finally, the skin was sutured, and animals were monitored daily after surgery. A buprenorphine injection was systematically given the next day.

To ensure that the effects observed after LPC injection were attributable to LPC and not to the surgical procedure itself, control injections with PBS were also performed (Figure S2). For these animals, the surgical procedure was identical to that described above.

#### rTSMS treatment

rTSMS was delivered with a commercially available figure of eight double coils featuring an aircooling system connected to a Magstim rapid² stimulator used for focal cortical and peripheral stimulations (Magstim, Whitland, UK). The coil was positioned in close contact with the back of the animal at the site of injury, using the mark located in the middle of the coil. The position was maintained using an articulated arm stand. According to manufacturer’s device manual, the size of the stimulated area was 1.5 cm².

rTSMS treatment was applied at a frequency of 10 Hz, 10 minutes per day during 7 days for *in vivo* experiments, and 3 days for *in vitro* ones. Stimulation protocol consisted of 10 s stimulation followed by 20 s of rest ^14,22^. For *in vivo* experiments, mice were kept under anesthesia with 2% of isoflurane during stimulation, the equivalent anesthesia was used for untreated animals. Peak magnetic intensity at the experimental distance was 0.4 T for *in vivo* experiments and 0.04 T for *in vitro* experiments.

#### Behavioral analyses

##### Open field test

Locomotor activity was assessed using the Open Field test. Mice were placed in a square open arena (50 × 50 cm) following a five-minute habituation period. Animals were then recorded for 3 minutes at 20 frames per second using a Basler acA2040-120μm camera. Video acquisition was performed with Pylon Viewer software. After recording, videos were processed using the tracking software DeepLabCut (DLC).

##### Video analyses

DLC was used for the functional analysis of locomotor behavior in mice. Developed in Python and based on deep learning techniques□^23^, DLC enables automated tracking of anatomical points of interest across large video datasets. After learning from manually annotated frames, the software generalizes point detection to new videos. In this study, as training, a total of 1.810 images were first annotated. For this, the default ResNet-50 neural network was used ^23^. The maximum number of training iterations was set to 200,000. The resulting model achieved an average error of 4.81 pixels and a test error of 5.51 pixels for images with a resolution of 2048 × 1536 pixels. Then, videos were analyzed using DLC. After processing, the software generated CSV files containing the tracked point coordinates. These data were then processed using custom Python scripts to calculate various kinematic parameters such as mean speed, total distance traveled, and immobility time. Because this unconstrained locomotor test is subject to high inter-individual variability and can be influenced by the animals’ motivation, we calculated a recovery index for these three parameters that accounts for this variability and evaluates functional recovery over time by comparing performances between D2 and D7, and between D2 and D14 for Figures 3 and S4 and between D2 and D9 for Figure 10. For these indices, a score of 0 indicates no improvement over time (motor abilities at D7, D14 or D9 being comparable to those at D0); a positive score reflects improved performance between D7, D14 or D9 relative to D2; and a negative score indicates a decline in performance over time. The recovery index for mean speed and distance traveled was calculated using the following formula: (D7 − D2)/D2, where the value obtained for each animal at the time of interest was used. The recovery index for immobility time was calculated according to the formula: (D2 − D7)/D2. For longer recovery periods, D7 was replaced by D14 or D9 as appropriate.

### Histological analyses

#### Tissue preparation and sectioning

An injection of a mixture of ketamine 1000 (0,75 µl/g) and xylazine (0,5 µl/g) was administered intraperitoneally to the mice. After verifying the absence of reflexes, a laparotomy and then a thoracotomy were performed to expose the heart. A needle was placed in the left ventricle. The right atrium was sectioned, 30 ml of 1X-EDTA PBS (1/100) was injected, and then 30 ml of 4% paraformaldehyde (antigenfix, f/P0016, MM france) was injected using a MasterFlex® peristaltic pump. The spinal cords were dissected, post-fixed overnight in 4% PFA and cryoprotected for ≥ 48 hours in 30 % sucrose (200-301-B, Euromedex). The tissues were embedded in Tissue-Tek OCT (Neg-50, 6502, Epedria), frozen in Isopentane between −30 °C and −40 °C, and then sectioned sagittally at 20 µm. The sections were mounted on slides and stored at −20 °C until use.

For the preparation of samples intended for RNASeq, the mice were anesthetized in the same manner as for the perfusions described above. After exposing the heart, a sterile catheter was placed in it and 25 mL of cold sterile 1X PBS was injected. After harvesting the spine, a 1.5 cm fragment was taken from the lesion site under sterile conditions on ice. The samples were instantly frozen in liquid nitrogen, and then stored at −80 °C until they were sent to the platform.

#### Immunohistochemistry

Identification of different cells and structures in spinal cord were conducted using primary antibodies. Samples were rehydrated with cold PBS. To reduce autofluorescence, a 0.1 M glycine bath was used for 20 min. The antibodies were diluted in 1X blocking buffer (ab126587, Abcam) and 0.5% Triton X-100 solution (93443, Sigma-Aldrich) to minimize non-specific binding. Antibodies incubation occurred overnight in a humidified chamber. The following primary antibodies were used: rabbit anti-platelet-derived growth factor α and β (PDGFR α/β, 1/500, Abcam, ab32570, RRID:AB_777165), mouse anti-glial fibrillary acidic protein (GFAP Cy3-conjugated, 1/500, Sigma-Aldrich, C9205, RRID:AB_476889), rabbit anti-ionized calcium-binding adapter molecule 1 (Iba1, 1/1000, SySy, 234-008, RRID:AB_2891288), rat anti-myelin basic protein (MBP, 1/100, Millipore, MAB 386, RRIB_94975).

After washing, antibody staining was detected with species-specific fluorescence-conjugated secondary antibodies (1/500, Jackson ImmunoResearch). Sections were counterstained with 4′,6-diamidino-2-phénylindole (DAPI; 1/5000, 1 μg/mL; Sigma-Aldrich) and mounted with Mowiol 4-88 reagent (475904, Sigma).

#### Image acquisition analysis and quantification

The slides were scanned using the automated Zeiss Axio Scan.Z1 system (epifluorescence microscopy) to digitize the immunohistochemical stainings. Three to five sections per animal were examined to identify the epicenter of the lesion, and an image of this epicenter was subsequently captured and analyzed. The stainings were then analyzed histologically and quantified using the QuPath® software (0.6.0 version). In these sagittal sections, negative GFAP-areas (astrocytic defect), positive PDGFrβ+ areas (fibrotic scar) and positive Iba1+ areas (inflammatory area) were measured. Additionally, three quantifications related to myelin were performed: the first measured the area lacking myelin (myelin defect), defined as the MBP-area; the second measured the total area containing myelin debris (Figure 1F); and the third quantified the total demyelinated area, corresponding to the sum of the two previous measurements.

### *In vitro* experiments

#### BV2 and RAW Cell line cultures

The BV2 line is an immortalized murine microglial cell line (ATCC CRL-2467, EOC) derived from the brain of a 10-day-old mouse. RAW 264.7 is an immortalized murine macrophage cell line (ATCC TIB-71) isolated from a mouse tumor that was induced by Abelson murine leukemia virus.

For both cell lines, the same culture protocol was used. Vials were thawed in a 37 °C water bath, and the cells were transferred to T25 flasks containing high-glucose DMEM (41965062, Gibco) supplemented with 10% fetal bovine serum heat inactivated (FBS, SH30088.03HI, Cytivia), 2% penicillin/streptomycin (15140122, Gibco) and 2 mM L-glutamine (25030081, Gibco). After 48 h, the cultures were first passaged: the medium was removed, cells were rinsed twice with PBS, exposed to 5 mL trypsin–EDTA (25300054, Gibco) for 3-5 min at 37 °C, and the reaction was quenched with complete medium. The suspension was centrifuged at 200g for 5 min, and the pellet was resuspended in 1 mL fresh medium. The cells were divided into four experimental conditions (Figures 7A and 8A)

⍰ BV2 or RAW control: cells were cultured during 5 days without specific treatment.
⍰ BV2 or RAW IL-1: At day 1, IL-1 (20 ng/ml) was added to the medium for 16 hours and then medium was removed and the cells were cultured for 4 days in standard medium.
⍰ BV2 or RAW + RMS: cells were treated with RMS for 3 days on days 2, 3 and 4.
⍰ BV2 or RAW IL-1+ RMS: cells were treated with IL-1 and RMS.

After 5 days in culture, the cells were harvested by trypsinization, pelleted, and snap-frozen at −80 °C until RNASeq analysis.

### RNA extraction, library construction, transcriptome sequencing and RNASeq analyses

The RNA from the spinal cord and cells was extracted, and sequenced by BGI Tech Solutions (Hong Kong, China).

#### Library Construction Methods

mRNA enrichment was performed on total RNA using oligo(dT)-attached magnetic beads. The enriched mRNA with poly(A) tails was fragmented using a fragmentation buffer, followed by reverse transcription using random N6 primers to synthesize cDNA double strands. The synthesized double-stranded DNA was then end-repaired and 5’-phosphorylated, with a protruding ‘A’ at the 3’end forming a blunt end, followed by ligation of a bubble-shaped adapter with a protruding ‘T’ at the 3’ end. The ligation products were PCR amplified using specific primers. The PCR products were denatured to single strands, and then single-stranded circular DNA libraries were generated using a bridged primer. The constructed libraries were quality-checked and sequenced after passing the quality control.

#### Sequencing

Sequencing was performed on DNBSEQ platform with PE150 (read length).

#### Analysis Pipeline

The sequencing data, also known as raw reads or raw data, were first subjected to quality control (QC) to determine whether they were suitable for further analysis. Once QC was completed, the clean reads were aligned to the reference sequences. Following alignment, a second QC was performed to assess the alignment quality, including alignment rate and the distribution of reads on the reference sequences. Subsequently, gene quantification analysis was carried out, along with various analyses based on gene expression levels (such as principal component analysis, correlation analysis, differential gene screening, etc.). Differentially expressed genes between samples were then investigated for gene ontology (GO) functional enrichment, pathway enrichment, clustering, protein-protein interaction networks, and transcription factor analysis.

#### Transcriptomic analyses

Transcriptomic analyses were conducted taking into account sex for *in vivo* samples, and experimental conditions for *in vitro* cell cultures. Male and female groups were analyzed separately. For each sex, differential gene expression analysis compared the LPC + rTSMS group with the LPC-only control group. Regarding immune cells, the macrophage RAW and microglial BV2 cell lines were analyzed independently. Several differential analyses were performed to compare experimental conditions. For each cell line, the following comparisons were conducted: 1) RMS vs CTRL, 2) CTRL vs CTRL + IL-1, 3) RMS + IL-1 vs CTRL + IL-1 and 4) RMS vs RMS + IL-1. These differential analyses allowed the identification of differentially expressed genes (DEGs), which were classified as either upregulated or downregulated.

Subsequently, GO enrichment analysis was performed to characterize the biological functions associated with the identified DEGs. GO terms are categorized into three main ontologies: biological process, molecular function, and cellular component. In the context of this study, biological processes were prioritized, as they are more likely to reveal key pathological mechanisms, such as inflammation or axonal demyelination. Analyses identified several dozen to hundreds of biological processes depending on the comparison. To better visualize potential interactions and relationships between biological processes, Cytoscape and GSEA software were used to model GO pathway enrichment networks[^24^. These networks were built using the results from differential expression analysis performed with the DESeq2 package[^25^. Integration into Cytoscape was done via the EnrichmentMap plugin, which allows visualization of enriched pathways as interconnected graphs based on gene overlap. To facilitate interpretation, the AutoAnnotate plugin was used to automatically cluster GO terms into functional thematic groups, using an algorithm based on semantic similarity.

Differential expression and enrichment analyses were performed using the DEVEA platform (v1.0). Normalized gene expression matrices were uploaded and analyzed according to the default pipeline, including identification of DEGs using DESeq2 and subsequent enrichment analysis. Pathway enrichment was conducted using GO and KEGG databases. Volcano plots, heatmaps, and dot plots were generated to visualize the most significantly dysregulated genes and pathways. Significance thresholds were set at adjusted p-value < 0.05 and log2 fold change > 1.

### Statistical analysis

All data are presented as mean ± standard deviation (SD). Statistical analyses were conducted using GraphPad Prism software, version for windows (GraphPad Software; 10.5.0). Before performing the statistical analyses proper, the normality of data distribution was assessed using the Shapiro–Wilk test. When the data followed a normal distribution, a parametric test (Student’s t-test) was applied; for datasets with a non-normal distribution, a non-parametric test (Mann–Whitney U test) was used. Detailed statistical analyses for each assay are provided in the figure legends. A p-value of less than 0.05 was considered statistically significant.

## RESULTS

### LPC injection induces a strong but transient inflammation and demyelination of the spinal cord, which is not observed following PBS injection

In the first part of this study, before investigating the effects of rTSMS, we aimed to determine the time course of LPC-induced demyelination, inflammation, and fibroglial scar formation in the ventral part of the spinal cord (Figure S1B-G). To this end, mice were injected with LPC and sacrificed for histological analyses at 8, 15, 22, and 29 days post-injection (Figure S1A). These analyses revealed that LPC induces robust demyelination and inflammation 8 days after injection, which already decreases at 15 days and nearly resolves by 22 and 29 days (Figure S1E-G). This process was also accompanied by the development of a fibrotic scar (PDGFrB-positive) and a local astrocytic defect (GFAP-negative) at the injection site, following a similar time course (Figure S1B-D). Although inflammation, demyelination, and fibroglial scar are still present 15 days after LPC injection, these parameters markedly decrease compared to day 8 (Figure S1B-G).

Next, we assessed the effects of LPC injection on locomotor functions over time using the open field test (Figure S1H-K). Computer-assisted analyses allowed us to quantify the mice mean speed, distance traveled, and immobility time; these parameters were compared before and after injection. The results showed that LPC injection significantly reduced the mean speed as early as day 2 post-injection (Figure S1I). This reduction persisted until day 4, after which it returned to baseline levels by day 7 (Figure S1I). Similarly, the distance traveled dropped at days 3 and 4 post-injection, but recovered to pre-injection values by day 7 (Figure S1J). In contrast, immobility time increased from day 2 post-injection and remained elevated until day 28 (Figure S1K). Together, these locomotor analyses demonstrate that LPC injection transiently impairs locomotor abilities, with a peak effect observed at day 3 post-injection (Figure S1I-K).

We then confirmed that inflammation, demyelination, and fibroglial scar formation were specifically due to LPC and not to the injection procedure itself. To this end, 10 female and 10 male mice were injected with PBS and perfused either at 8 or 15 days post-injection for histological analyses, and compared to LPC-injected mice (Figure S2A-N). Unlike LPC, PBS injection did not induce demyelination, inflammation, or fibroglial scar formation either in male or female mice (Figure S2B-K), confirming that the observed effects were specifically due to LPC.

Similarly, we verified that PBS injection had no impact on locomotor function (Figure S2A and L-N). As described above, we analyzed average velocity, distance traveled, and immobility time in PBS-injected mice and compared them to LPC-injected animals. These analyses confirmed that PBS injection had no effect on locomotor parameters at any of the analyzed time points (Figure S2L-N).

### When applied at the presymptomatic stage, rTSMS reduces demyelination and inflammation and modulates the fibroglial scar in LPC model

In a first set of experiments, we investigated the effects of rTSMS in both female and male mice. These experiments were carried out up to 15 days after LPC injection, based on the results shown in Figure S1 indicating that LPC injection has limited effects beyond this time point (Figure S1). All animals received LPC injections, and half of them were subjected to rTSMS treatment for 7 consecutive days, starting one day after LPC injection (Figure 1B). Mice were perfused at day 8 post-LPC for histological analyses (Figure 1B). Our results show that rTSMS markedly reduced inflammation and the total demyelinated area in this experimental paradigm (Figure 1C–E). The total demyelinated area actually represents both the region devoid of myelin and the area where myelin is degraded and present in the form of debris. Analyses of these two complementary processes have been performed and revealed that rTSMS significantly reduced both processes 8 days after LPC injection (Figure 1G and H).

These experiments were repeated in independent cohorts to investigate the effects of rTSMS when mice were perfused 15 days after LPC injection, with the rTSMS protocol starting on day 1 and lasting for 7 days (Figure 1I). At this later time point, rTSMS no longer exerted significant effects on inflammation or overall demyelination (Figure 1J–L), nor on demyelinated areas or myelin debris within the spinal parenchyma (Figure 1M and N).

It has been demonstrated that fibrosis within spinal and brain tissue is not only observed in traumatic pathologies but also in inflammatory demyelinating diseases ^26^. Therefore, we investigated the effects of rTSMS on the formation of the fibroglial scar in this LPC injection model at 8 and 15 days post-injection (Figure 2A and E). Our results show that rTSMS reduced fibrosis and also attenuated the astrocytic defect within the lesion area at 8 days after LPC injection (Figure 2B–D).

**Figure 2:**
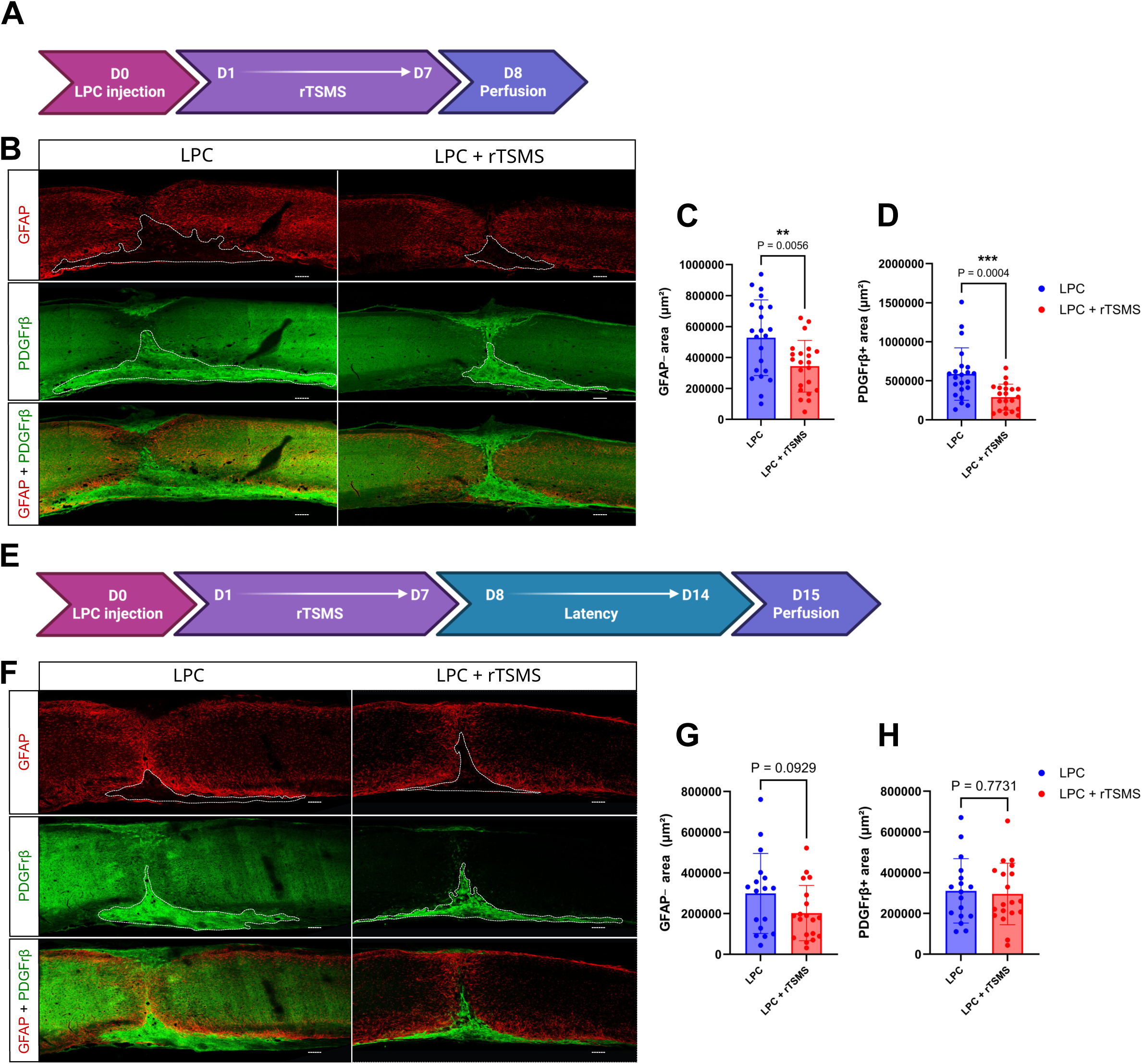
rTSMS modulates the fibroglial scar in LPC model, when applied at the presymptomatic stage. **A.** Experimental design: on day 0, all mice received an LPC injection; half of them underwent rTSMS treatment from D1 to D7, and all mice were perfused at D8 for histological analyses. **B-D.** Analysis of the effects of rTSMS on fibroglial scar formation 8 days after LPC injection with LPC group in blue and LPC + rTSMS group in red. **B.** Representative pictures of sagittal spinal cord sections of LPC and LPC + rTSMS groups 8 days after LPC injection. Sections were stained with anti-GFAP (in red) and anti-PDGFrβ (in green) antibodies. The scale bar represents 200µm **C**. Quantification of GFAP-area. **D.** Quantification of PDGFrβ+ area. **E.** Experimental design: on day 0, all mice received an LPC injection; half of them underwent rTSMS treatment from D1 to D7, and all mice were perfused at D15 for histological analyses. **F-H.** Analysis of the effects of rTSMS on fibroglial scar formation 15 days after LPC injection with LPC group in blue and LPC + rTSMS group in red. **F.** Representative pictures of sagittal spinal cord sections of LPC and LPC + rTSMS groups 15 days after LPC injection. Sections were stained with anti-GFAP (in red) and anti-PDGFrβ (in green) antibodies. The scale bar represents 200µm **G**. Quantification of GFAP-area. **H.** Quantification of PDGFrβ+ area. N = 17-22 mice in the LPC group and N = 17-22 mice in the LPC + rTSMS group. Statistical analyses were performed using Unpaired t-test (C, G and H) and Mann-Whitney test (D) (**=P≤ 0.01 and ***=P≤ 0.001).

Similarly to what was observed for inflammation and demyelination (Figure 1), our data demonstrate that at 15 days post-LPC injection, rTSMS no longer exerted a significant effect on fibroglial scar (Figure 2F–H), although a trend toward reduced GFAP negative area was still detected in the rTSMS group (Figure 2G).

Since inflammatory diseases such as MS display a strong sex effect ^27^, we investigated whether LPC injection or subsequent rTSMS treatment differentially affected tissue responses in male versus female mice (Figure S3A–N). At 8 days post-LPC injection, both male and female mice exhibited comparable fibroglial scar and inflammation (Figure S3B–D), as well as similar total demyelinated areas and myelin debris (Figure S3E and F). Interestingly, the demyelinated area (MBP defect) was smaller in males compared to females at this time point (Figure S3G). Likewise, both sexes exhibited comparable responses to rTSMS treatment for all these parameters at day 8 post-LPC (Figure S3B–G). At 15 days post-LPC injection, male and female mice again displayed similar fibroglial scar, inflammation and demyelination, regardless of whether they had received LPC alone or LPC combined with rTSMS (Figure S3H–N).

Our results show that initiating rTSMS the day after LPC injection reduces inflammation and demyelination, and modulates the fibroglial scar, with comparable effects in male and female mice.

### When applied at the presymptomatic stage, rTSMS enhances locomotor functions in LPC model

In order to correlate these histological findings with functional outcomes, we performed locomotor behavioral analyses by monitoring mice movements before LPC injection, and then from the second day post-injection up to day 14 using the open field test (Figure 3A). Our results show that rTSMS does not modulate the average locomotor speed over time (Figure 3B). Indeed, no significant difference was observed between the LPC and rTSMS groups for this parameter (Figure 3B). In contrast, rTSMS increases the total distance traveled and decreases immobility time (Figure 3C and D, respectively). Notably, mice receiving rTSMS displayed an improvement in these parameters after the treatment was discontinued, with enhanced distance traveled at day 14 and reduced immobility time at days 11 and 14 (Figure 3C and D, respectively).

**Figure 3:**
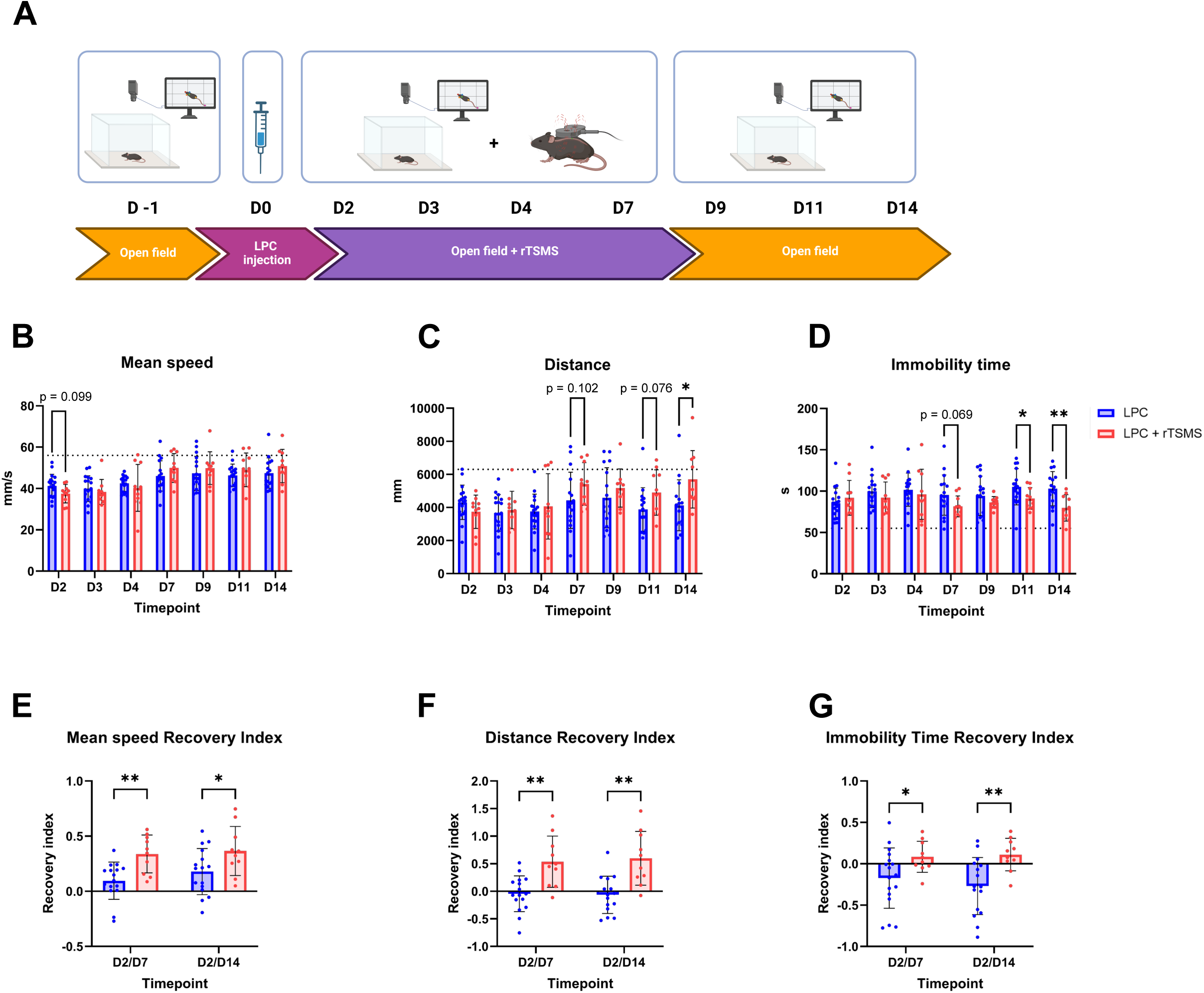
rTSMS enhances locomotor functions in LPC model, when applied at the presymptomatic stage. **A.** Experimental design: on day 0, all mice received an LPC injection; half of them underwent rTSMS treatment from D1 to D7, in parallel locomotor behaviors were recorded for all the mice one day before surgery and at D2, D3, D4, D7, D9, D11 and D14 after LPC injection. **B-D.** Analysis of the effects of rTSMS on functional recovery in a free-field locomotion open field test analysis over time (from 2 to 14 days post-LPC injection) with LPC mice in blue and LPC + rTSMS mice in red. **B.** Quantification of mean speed. **C.** Quantification of distance traveled. **D.** Quantification of immobility time. **E-G.** Analysis of the effects of rTSMS on recovery indices with LPC mice in blue and LPC + rTSMS mice in red. Recovery indices were calculated for each animal by comparing the values measured at day 2 (D2) to those measured at day 7 (D7) and day 14 (D14) post-lesion. **E.** Recovery indice for mean speed. **F.** Recovery indice for distance traveled. **G.** Recovery indice for immobility time. N = 16 mice in the LPC group and N = 10 mice in the LPC + rTSMS group. The dotted line represents pre-lesion values. Statistical analyses were performed using Welch’s t-test (*=P≤ 0.05; **=P≤ 0.01).

Because this unconstrained locomotor test is subject to high inter-individual variability and can be influenced by the animals’ motivation, we calculated a recovery index for these three parameters that accounts for this variability and evaluates functional recovery over time by comparing performances between D2 and D7, and between D2 and D14 (Figure 3E–G). As described above, for these indices, a score of 0 indicates no improvement over time (motor abilities at D7 or D14 similar to those at D0); a positive score reflects improved performance between D7 and D14 compared to D2; and a negative score indicates a decline in performance. Analysis of these indices revealed that, for the three parameters, rTSMS-treated animals showed greater motor recovery compared to controls, both at 7 and 14 days post-injection (Figure 3E–G).

As done previously, we sought to verify that no treatment effect could be attributed to sex differences. We therefore compared LPC and rTSMS groups by analyzing males and females separately (Figure S4). We found that rTSMS induced an early restoration of locomotor functions in males, with significant improvements in average speed and all three recovery indices at day 7 post-LPC injection (Figure S4G-L). In females, however, rTSMS effects appeared later, at day 14, particularly improving distance traveled, immobility time, and the corresponding recovery indices (Figure S4A-F).

Altogether, our results demonstrate that initiating rTSMS the day after LPC injection, rTSMS enhances locomotor performance in both male and female mice.

### rTSMS modulates immune response with a strong sex-dependent effect

In order to better characterize the pleiotropic effects of rTSMS in this inflammatory model, we performed RNASeq experiments separately in female and male mice, comparing LPC and LPC + rTSMS groups (Figures 4 and 5, respectively). All animals received LPC injections, and half of them were subjected to rTSMS treatment for 7 consecutive days, starting one day after LPC injection, then all mice were euthanized on D8 to perform RNASeq experiments (Figure 4A and 5A). These analyses identified a large number of biological processes that were dysregulated between LPC and LPC + rTSMS animals in both sexes (Figures 4 and 5). Notably, the number of dysregulated processes was higher in females compared to males (Figures 4 and 5, respectively). Most of these processes were associated with immune and inflammatory responses, cell death, neuronal plasticity, cellular respiration, as well as general cellular and metabolic functions (Figures 4 and 5). A clear sex-dependent difference was observed in the distribution of these functional categories. In females, immune, inflammatory, and cell death processes were enriched in the LPC group, whereas neuronal processes were enriched in the LPC + rTSMS group (Figure 4). Conversely, in males, immune and inflammatory processes were predominantly found in the LPC + rTSMS group, whereas neuronal processes were more strongly represented in the LPC group (Figure 5). Moreover, processes related to the innate response are also present in males (Figure 5). This inversion suggests a differential treatment response between sexes, both at the immune and neuronal levels. Moreover, females displayed a greater number of altered immune and inflammatory processes, suggesting increased sensitivity and a sex-dependent effect of rTSMS on the modulation of these pathways (Figure 4).

**Figure 4:**
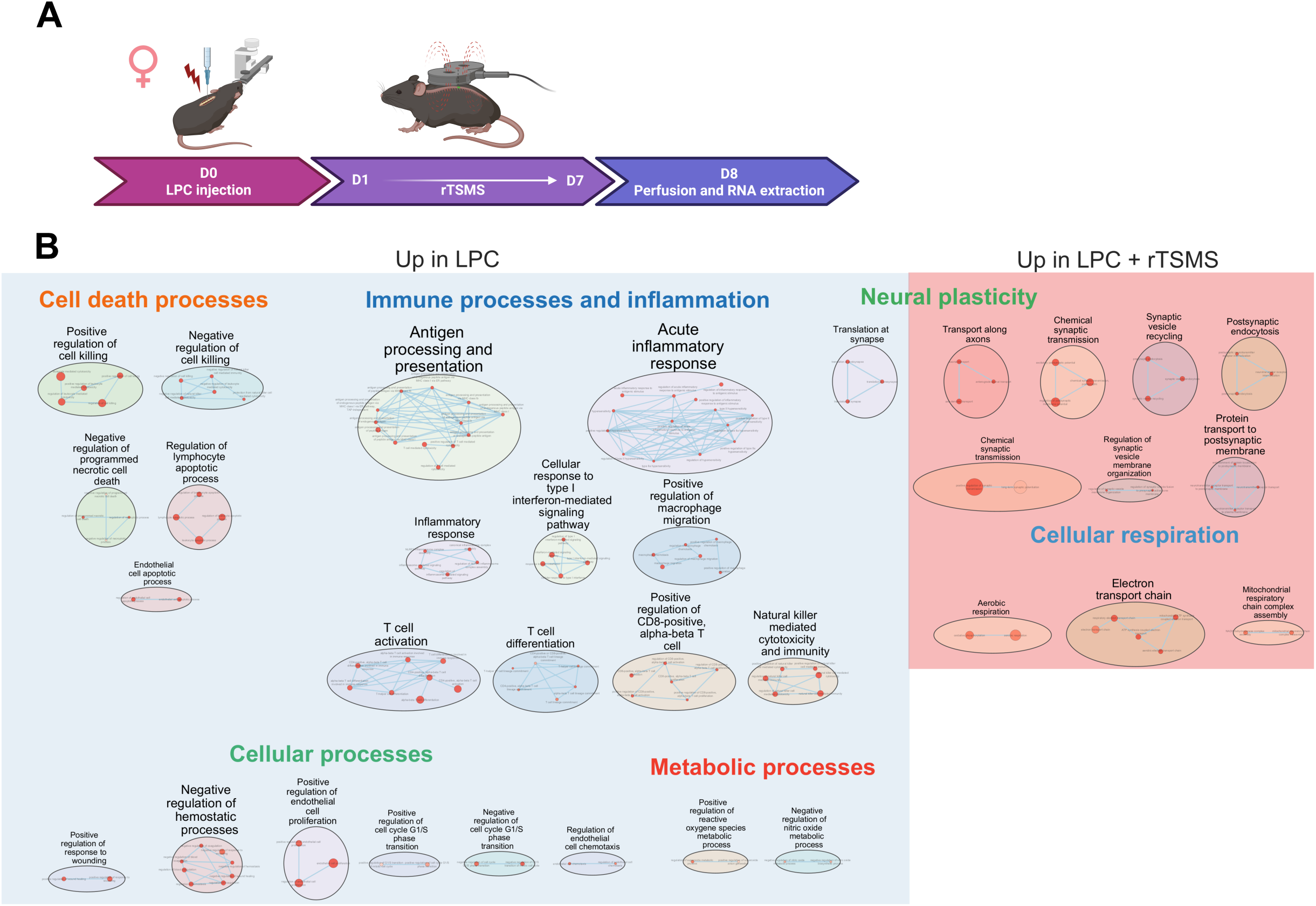
rTSMS modulates gene expression in female mice. **A.** Experimental design: on day 0, all female mice received an LPC injection; half of them underwent rTSMS treatment from D1 to D7 and all mice were perfused at D8 for RNASeq experiments. **B.** Cytoscape representation of deregulated GO terms (“Biological Process” category). Each dot represents a deregulated GO-Term; their size and color are, respectively, proportional to the number of genes in the GO-term and the enrichment adjusted p-value (false discovery rate q value). Blue indicates GO terms upregulated in the LPC group, while red indicates those upregulated in the LPC + rTSMS group. GO-Terms are grouped into categories using the Autoannotate Cytoscape app (plugin). N = 6 mice in the LPC group and N = 6 mice in the LPC + rTSMS group.

**Figure 5:**
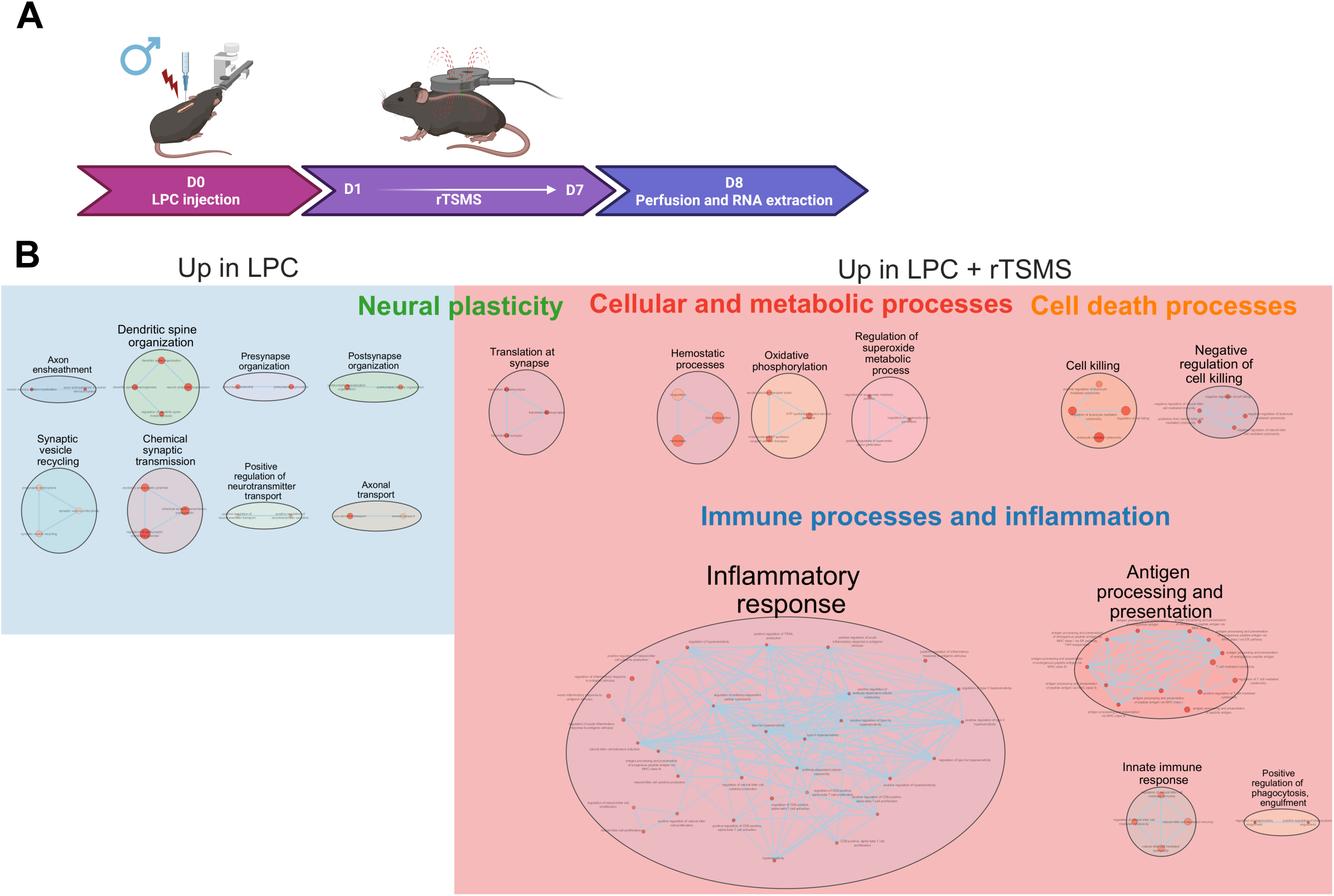
rTSMS modulates gene expression in male mice. **A.** Experimental design: on day 0, all male mice received an LPC injection; half of them underwent rTSMS treatment from D1 to D7 and all mice were perfused at D8 for RNASeq experiments. **B.** Cytoscape representation of deregulated GO terms (“Biological Process” category). Each dot represents a deregulated GO-Term; their size and color are, respectively, proportional to the number of genes in the GO-term and the enrichment adjusted p-value (false discovery rate q value). Blue indicates GO terms upregulated in the LPC group, while red indicates those upregulated in the LPC + rTSMS group. GO-Terms are grouped into categories using the Autoannotate Cytoscape app (plugin). N = 6 mice in the LPC group and N = 6 mice in the LPC + rTSMS group.

To gain a functional perspective by linking the differentially regulated biological functions between LPC and rTSMS groups in males and females, we conducted GSEA analyses (Figure 6). These analyses revealed that NF-κB signaling, neutrophil extracellular trap formation, IL-17 signaling, Th17 cell differentiation, and TNF signaling pathways were downregulated by rTSMS in females (Figure 6A), highlighting the regulatory role of this treatment on inflammatory and adaptive immune pathways in females. In contrast, in male mice, there is an upregulation of lysosome, phagosome, TGF-β signaling, neutrophil extracellular trap formation, and leukocyte transendothelial migration pathways in the stimulated group compared to controls (Figure 6B), indicating that rTSMS primarily regulates phagocytosis and innate immunity in males. Altogether, these findings highlight the strong impact of rTSMS on immune response regulation and reveal a pronounced sex-dependent effect of this treatment.

**Figure 6:**
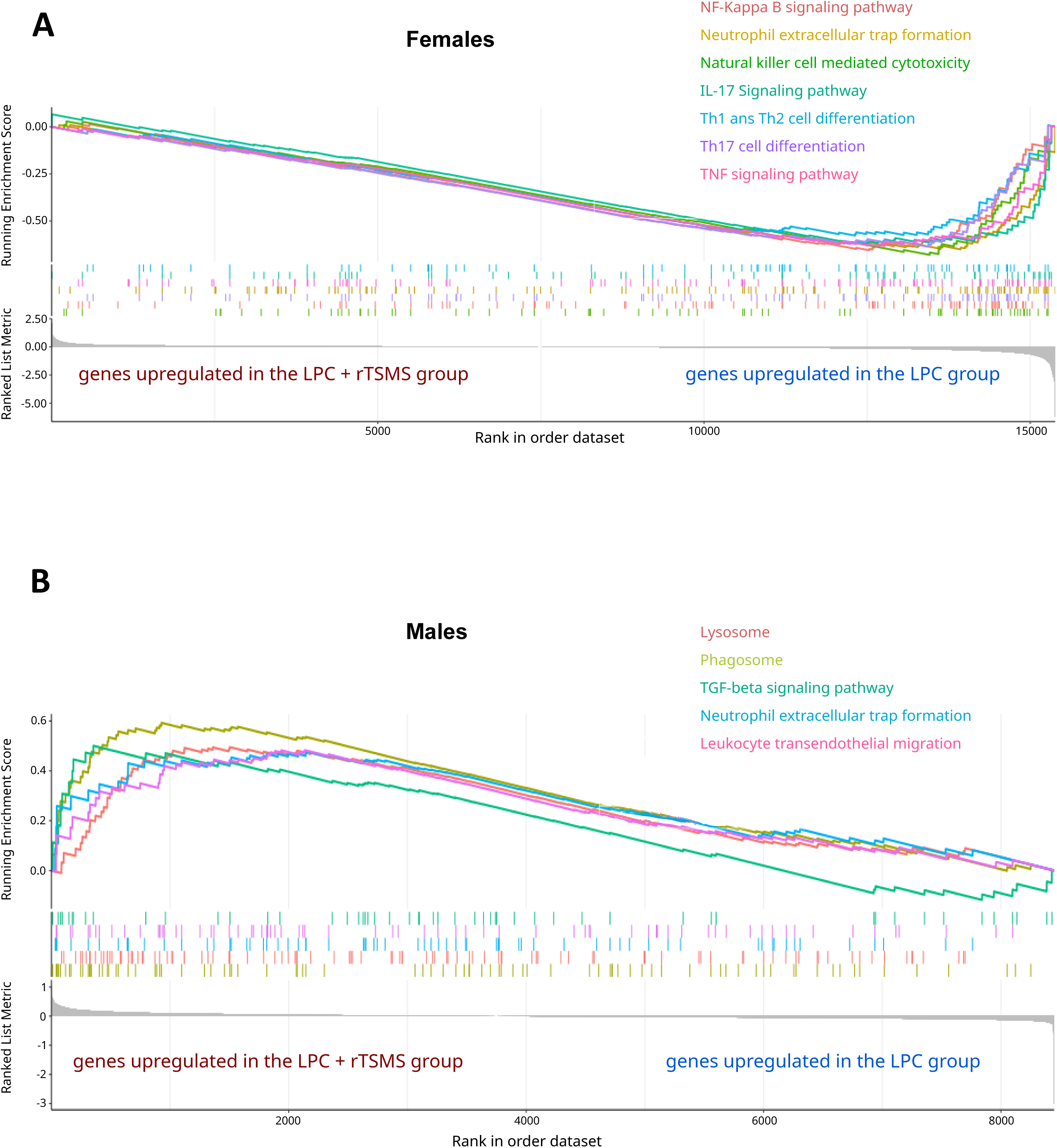
rTSMS modulates differential gene expression in male and female mice 8 days after LPC injection. GSEA (Gene Set Enrichment Analysis) identifies enriched biological pathways within a set of genes ranked according to their differential expression levels after rTSMS and LPC 8 days post-LPC injection in female (**A**) and male (**B**) mice.

### RMS differentially modulates microglial and macrophage responses to IL-1 and attenuates pro-inflammatory signaling

Since RNASeq analyses revealed that rTSMS strongly modulates immune and inflammatory processes, we next investigated the effects of RMS *in vitro* in microglial (BV2) and macrophage (RAW) cell lines. In a first series of experiments, half of these cell cultures were exposed to RMS for 3 days and collected the following day for RNASeq experiments (Figure 7A and 8A left panels). Then, in a second step, the cells were activated by the addition of IL-1, a pro-inflammatory cytokine, to the culture medium for 16 hours, after which half of them received RMS treatment (Figure 7A and 8A right panels) ^28^. We first examined the effects of RMS on the IL-1-unactivated BV2 microglial cell line (Figure 7B). Comparison between RMS and CTRL conditions showed that RMS significantly altered the transcriptomic profiles of BV2 cells (Figure 7B). Specifically, RMS induced enrichment of biological processes related to immune processes and inflammation such as Natural Killer cells and T lymphocytes, neuronal plasticity, and endothelial cells, suggesting a modulation of immune and neuroinflammatory responses in this cell line (Figure 7B).

**Figure 7:**
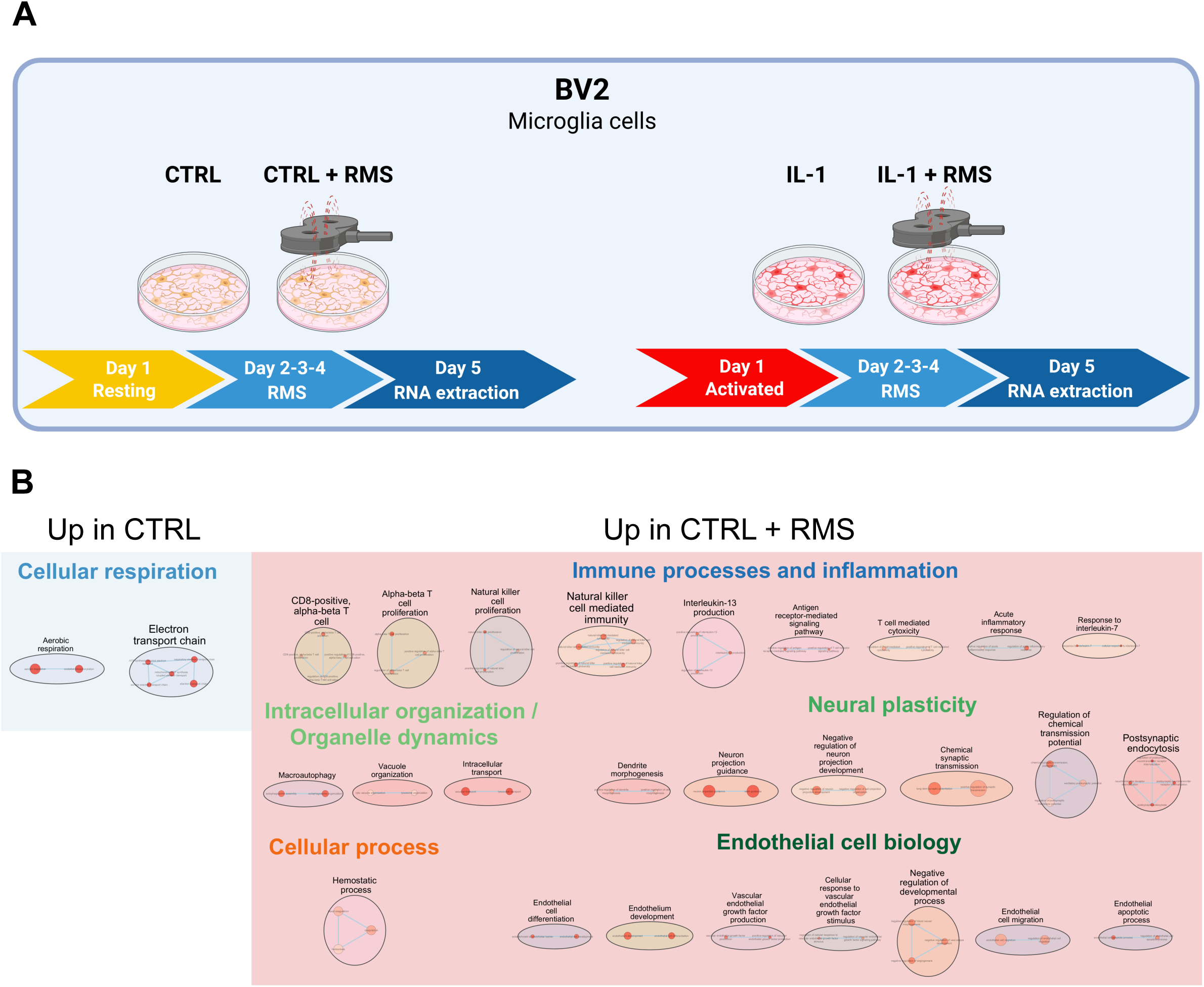
RMS modulates gene expression in BV2 microglial cell cultures when not activated by IL-1. **A.** Experimental design: BV2 cells were divided into four experimental conditions according to the arrangement of the culture plates from left to right: control (CTRL), control treated with RMS (CTRL + RMS), IL-1 activated cells (IL-1), and IL-1 activated cells treated with RMS (IL-1 + RMS). (1) For the non-activated cells, cultures were initiated on day 1; half of them received RMS treatment on days 2, 3 and 4, and all cultures were subsequently harvested for RNASeq experiments on day 5. (2) For IL-1 activated paradigm IL-1 was added on day 1. Then, RMS treatment was applied daily on days 2, 3, and 4 and RNA was extracted on day 5. **B.** Cytoscape representation of deregulated GO terms (“Biological Process” category). Each dot represents a deregulated GO-Term; their size and color are, respectively, proportional to the number of genes in the GO-term and the enrichment adjusted p-value (false discovery rate q value). Blue indicates GO terms upregulated in the BV2 group, while red indicates those upregulated in the CTRL + RMS group. GO-Terms are grouped into categories using the Autoannotate Cytoscape app (plugin). N = 5 independents samples in the CTRL group and N = 5 independents samples in the CTRL + RMS group.

**Figure 8:**
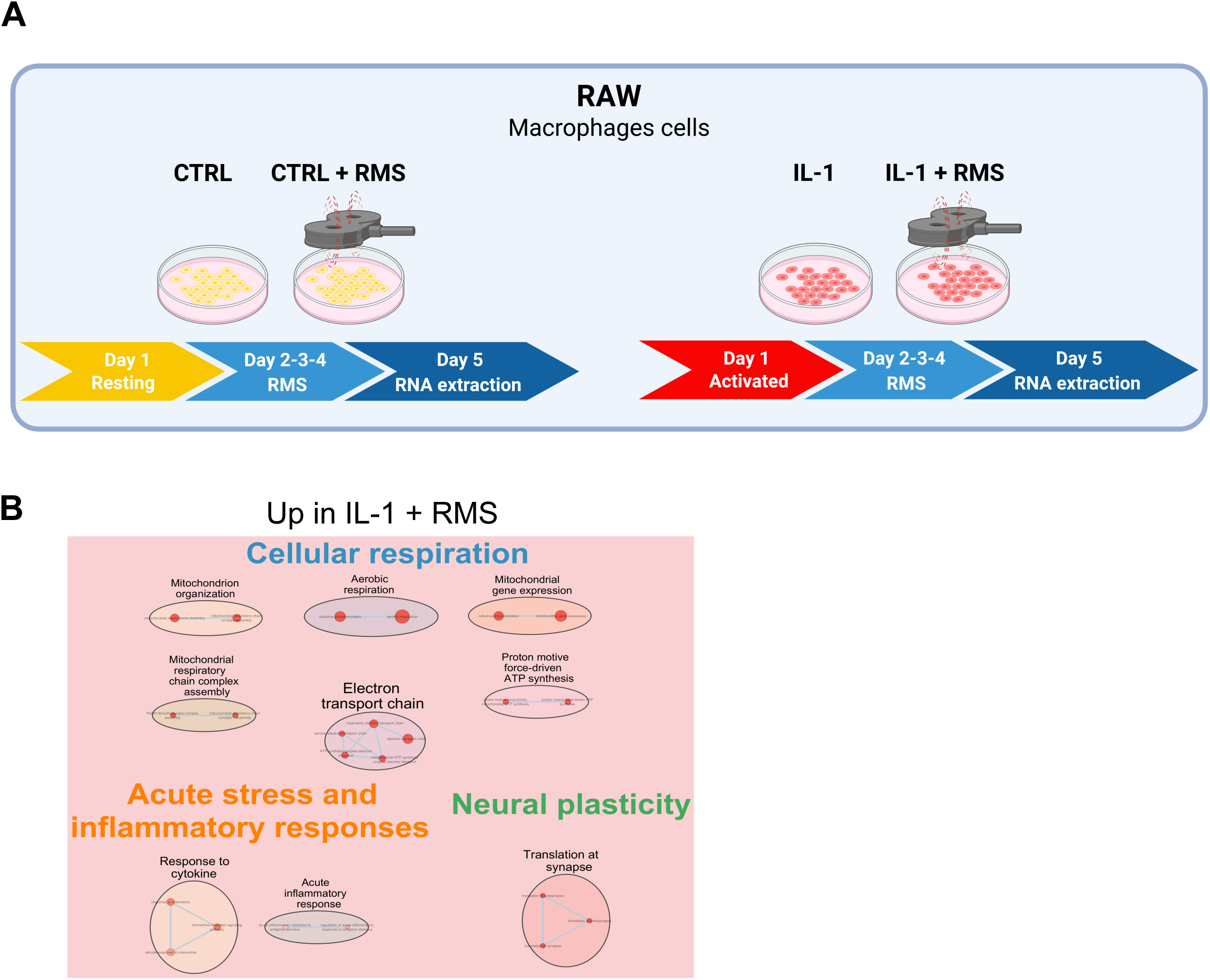
RMS modulates gene expression in RAW macrophage cell cultures when activated by IL-1. **A.** Experimental design: RAW cells were divided into four experimental conditions according to the arrangement of the culture plates from left to right: control (CTRL), control treated with RMS (CTRL + RMS), IL-1 activated cells (IL-1), and IL-1 activated cells treated with RMS (IL-1 + RMS). (1) For the non-activated cells, cultures were initiated on day 1; half of them received RMS treatment on days 2, 3 and 4, and all cultures were subsequently harvested for RNASeq experiments on day 5. (2) For IL-1 activated paradigm IL-1 was added on day 1. Then, RMS treatment was applied daily on days 2, 3, and 4 and RNA was extracted on day 5. **B.** Cytoscape representation of deregulated GO terms (“Biological Process” category). Each dot represents a deregulated GO-Term; their size and color are, respectively, proportional to the number of genes in the GO-term and the enrichment adjusted p-value (false discovery rate q value). Red indicates GO terms upregulated in the CTRL + RMS group. GO-Terms are grouped into categories using the Autoannotate Cytoscape app (plugin). N = 5 independents samples in the IL-1 group and N = 5 independents samples in the IL-1 + RMS group.

We then reproduced these experiments in the macrophage RAW cell line, and found that, unlike BV2 microglial cells, RMS had no effect under non-activated conditions.

Subsequently, we investigated the effects of RMS in BV2 and RAW cell lines after activation with the pro-inflammatory cytokine IL-1 for 16 hours (Figure 7A and 8A right panels). Before assessing RMS effects under these conditions, RNASeq analyses confirmed IL-1–induced activation, with a strong dysregulation of immune and inflammatory processes in BV2 + IL-1 compared to control BV2 cells (Figure S5). Similar IL-1–driven activation was observed in RAW cells (Figure S6). We then assessed the effects of RMS in IL-1–activated cells by comparing BV2/RAW + IL-1 and BV2/RAW + IL-1 + RMS groups. Under these conditions, RMS had no detectable effect on gene expression in BV2 cells. By contrast, RMS modulated processes related to cellular respiration, inflammation, and neuronal plasticity in RAW cells (Figure 8B).

Finally, we sought to evaluate whether RMS could counteract the effects of IL-1. As described above, IL-1 strongly impacts BV2 and RAW cells by inducing a large number of immune and inflammatory processes (Figures S5 and S6). We therefore compared BV2/RAW + RMS versus BV2/RAW + IL-1 + RMS groups. Strikingly, under these conditions, RMS markedly reduced the number of IL-1–induced dysregulated processes in BV2 cells (Figure S7 to compare to Figure S5) and completely abolished IL-1 effects in RAW cells.

Altogether, our results show that RMS: 1. modulates the transcriptional profile of microglia when not activated by IL-1; 2. modulates the transcriptional profile of macrophages when activated by IL-1 and; 3. attenuates IL-1 effects in macrophages and fully blocks them in microglia.

### When applied at the symptomatic stage, rTSMS reduces demyelination and inflammation and modulates the fibroglial scar in LPC model only in female mice

We next focused on a more pre-clinical paradigm in which rTSMS was initiated after the onset of motor deficits, a time window that could correspond in Humans to the point of medical intervention. To this end, treatment was started 3 days after LPC injection, the time point when motor impairments are most pronounced (Figure S1I-K). As in previous experiments, mice were injected on day 0 and received rTSMS treatment for 7 days, here from day 3 to day 9, and were perfused on day 10 for histological analyses (Figure 9A).

**Figure 9:**
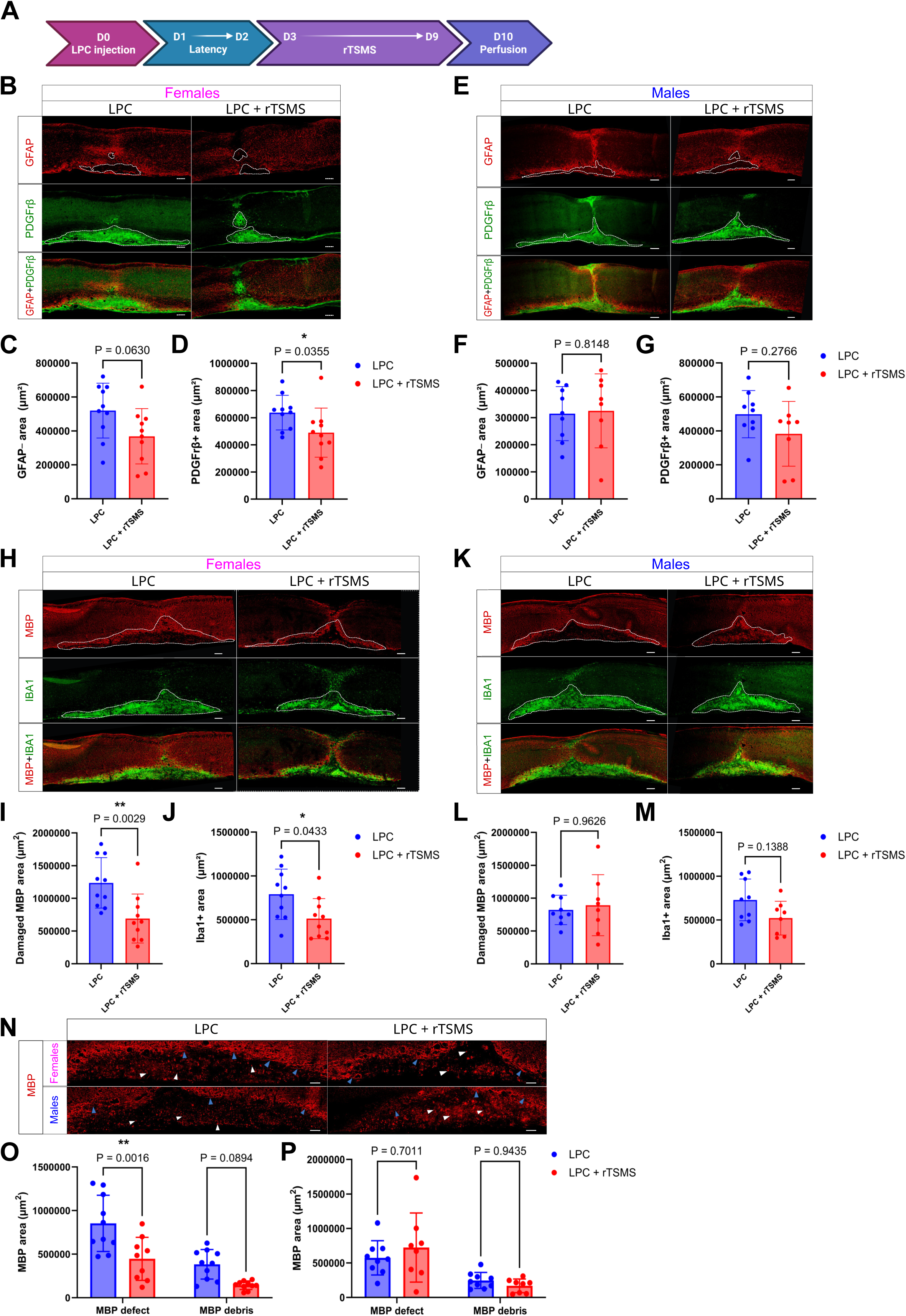
rTSMS reduces demyelination and inflammation and modulates the fibroglial scar in LPC model only in female mice, when applied at the symptomatic stage. **A.** Experimental design: on day 0, all mice received an LPC injection; half of them underwent rTSMS treatment from D3 to D9 (during 7 consecutive days), and all mice were perfused at D10 for histological analyses. **B-D.** Analysis of the effects of rTSMS on fibroglial scar formation 10 days after LPC injection in female mice with LPC group in blue and LPC + rTSMS group in red. **B.** Representative pictures of sagittal spinal cord sections of LPC and LPC + rTSMS groups 10 days after LPC injection. Sections were stained with anti-GFAP (in red) and anti-PDGFrβ (in green) antibodies. The scale bar represents 200 µm **C**. Quantification of GFAP-area. **D.** Quantification of PDGFrβ+ area. **E-G.** Analysis of the effects of rTSMS on fibroglial scar formation 10 days after LPC injection in male mice with LPC group in blue and LPC + rTSMS group in red. **E.** Representative pictures of sagittal spinal cord sections of LPC and LPC + rTSMS groups 10 days after LPC injection. Sections were stained with anti-GFAP (in red) and anti-PDGFrβ (in green) antibodies. The scale bar represents 200 µm **F**. Quantification of GFAP-area. **G.** Quantification of PDGFrβ+ area. **H-J.** Analysis of the effects of rTSMS on inflammation and damaged MBP area 10 days after LPC injection in female mice with LPC group in blue and LPC + rTSMS group in red. **H.** Representative pictures of sagittal spinal cord sections of LPC and LPC + rTSMS groups 10 days after LPC injection. Sections were stained with anti-MBP (in red) and anti-Iba1 (in green) antibodies for Iba1 and damaged MBP area analyses. The scale bar represents 200 µm. **I**. Quantification of Iba1+ area. **J.** Quantification of damaged MBP area (MBP defect + MBP debris areas). **K-M.** Analysis of the effects of rTSMS on inflammation and damaged MBP area 10 days after LPC injection in male mice with LPC group in blue and LPC + rTSMS group in red. **K.** Representative pictures of sagittal spinal cord sections of LPC and LPC + rTSMS groups 10 days after LPC injection. Sections were stained with anti-MBP (in red) and anti-Iba1 (in green) antibodies for Iba1 and damaged MBP area analyses. The scale bar represents 200 µm. **L**. Quantification of damaged MBP area (MBP defect + MBP debris areas) **M.** Quantification of Iba1+ area. **N-P.** Analysis of the effects of rTSMS on demyelination processes 10 days after LPC injection in female and male mice with LPC group in blue and LPC + rTSMS group in red. **N.** Representative pictures of sagittal spinal cord sections of LPC and LPC + rTSMS groups 10 days after LPC injection. Sections were stained with anti-MBP (in red) antibody to analyse MBP defect and MBP debris areas. The white arrows indicate myelin debris, and the blue arrows indicate axons undergoing demyelination. **O**. Quantification of MBP defect (MBP-area) and quantification of MBP debris area in females with LPC group in blue and LPC + rTSMS group in red. **P.** Quantification of MBP defect (MBP-area) and quantification of MBP debris area in males with LPC group in blue and LPC + rTSMS group in red. N = 10 mice in the LPC group for female mice and N= 9 mice in the LPC group for male mice; N = 10 mice in the LPC + rTSMS group for female mice and N = 8 in the LPC + rTSMS group for male mice. Statistical analyses were performed using Mann-Whitney t-test (*=P≤ 0.05; **=P≤ 0.01).

In this paradigm, a sex-dependent effect of treatment was observed, prompting us to analyze male and female mice separately. Results showed that in females, rTSMS tended to reduce astrocytic defects (Figure 9B and C) and significantly decreased fibrosis, inflammation, and demyelination (Figure 9B, D, H, I, J, N, and O). In contrast, rTSMS had no effect in males, regardless of the parameter measured (Figure 9E, F, G, K, L, M, N, and P). These findings demonstrate that, when applied at the symptomatic stage, rTSMS reduces demyelination and inflammation and modulates the fibroglial scar in the LPC model, but only in female mice.

We then assessed the impact of this treatment on locomotor functions through behavioral analyses (Figure 10). Locomotor performances were assessed before LPC injections and at days 2, 3, 4, 7 and 9 post-injection (Figure 10A). In this paradigm, recovery indices were calculated by comparing locomotor parameters between D2 and D9, in correlation with the rTSMS treatment period. Again, males and females were analyzed separately due to the marked sex effect. In females, rTSMS induced a moderate motor recovery over time, with improved distance traveled and reduced immobility time at day 7 (Figures 10C and D), without effects on average speed or recovery indices (Figures 10B and E–G respectively). In males, analyses revealed group differences in all three locomotor parameters as early as day 2—i.e., prior to treatment onset—suggesting stronger baseline inter-group variability than in previous experiments (Figures 10H–J). In this context, the recovery indices proved particularly informative (Figures 10K–M). These indices showed that rTSMS did not improve motor recovery in males and even tended to reduce the recovery index for distance traveled in the treated group (Figures 10K–M). Altogether, our results demonstrate that when applied at the symptomatic stage, rTSMS induces moderate functional recovery in the LPC model, but only in female mice.

**Figure 10:**
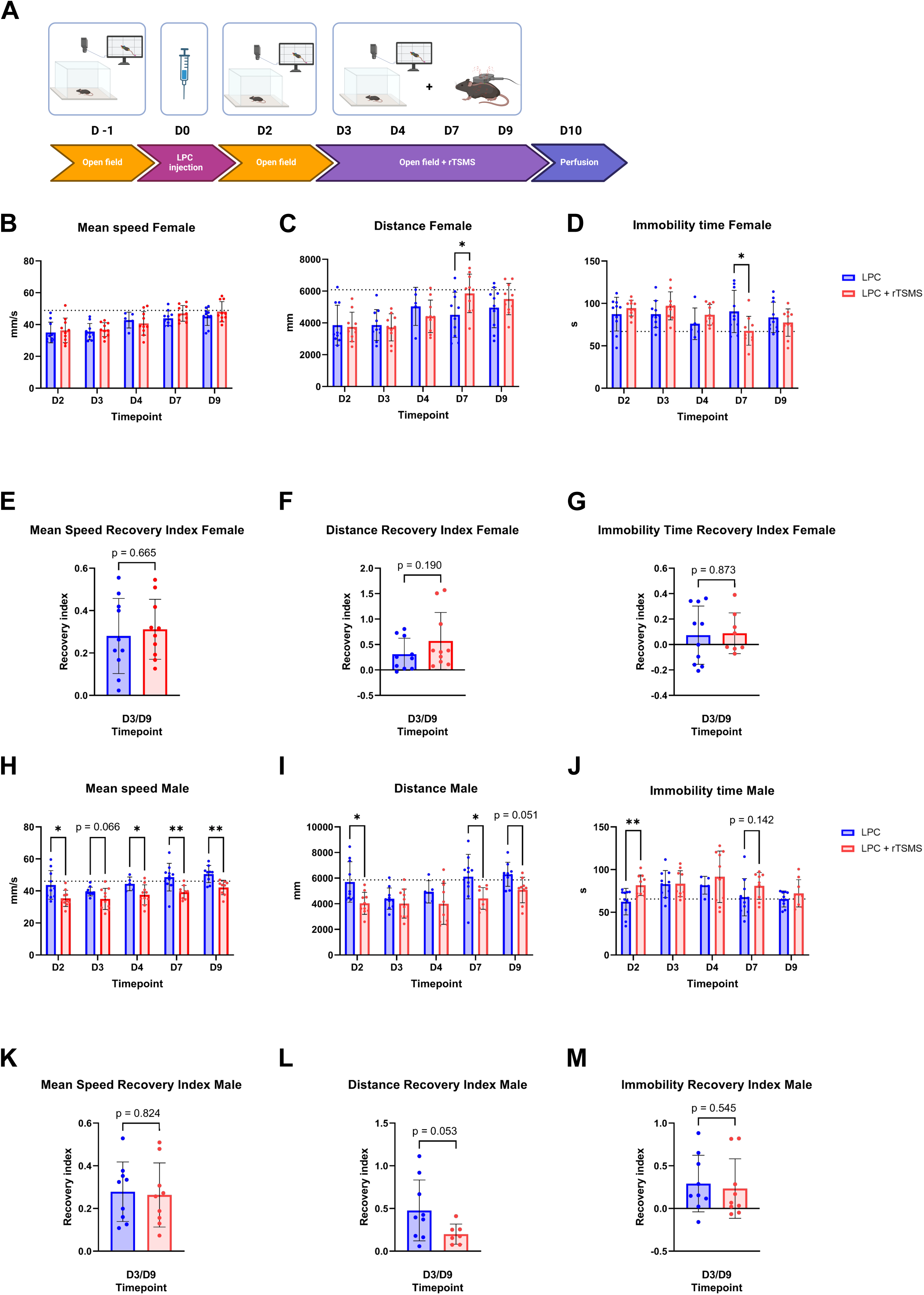
rTSMS induces moderate functional recovery in female mice in LPC model when applied at the symptomatic stage, but fails to do so in males. **A.** Experimental design: on day 0, all mice received an LPC injection; half of them underwent rTSMS treatment from D3 to D9 (during 7 consecutive days), in parallel locomotor behaviors were recorded for all the mice one day before surgery and at D2, D3, D4, D7 and D9 after LPC injection. **B-D.** Analysis of the effects of rTSMS on functional recovery in a free-field locomotion - open field test analysis over time (from 2 to 9 days post-LPC injection) with LPC mice in blue and LPC + rTSMS mice in red in female mice. **B.** Quantification of mean speed. **C.** Quantification of distance traveled. **D.** Quantification of immobility time. **E-G.** Analysis of the effects of rTSMS on recovery indices with LPC mice in blue and LPC + rTSMS mice in red. Recovery indices were calculated for each animal by comparing the values measured at day 2 (D2) to those measured at day 9 (D9) post-lesion in female mice. **E.** Recovery indice for mean speed. **F.** Recovery indice for distance traveled. **G.** Recovery indice for immobility time. **H-J.** Analysis of the effects of rTSMS on functional recovery in a free-field locomotion - open field test analysis over time (from 2 to 9 days post-LPC injection) with LPC mice in blue and LPC + rTSMS mice in red in male mice. **H.** Quantification of mean speed. **I.** Quantification of distance traveled. **J.** Quantification of immobility time. **K-M.** Analysis of the effects of rTSMS on recovery indices with LPC mice in blue and LPC + rTSMS mice in red in male mice. Recovery indices were calculated for each animal by comparing the values measured at day 2 (D2) to those measured at day 9 (D9) post-lesion. **K.** Recovery indice for mean speed. **L.** Recovery indice for distance traveled. **M.** Recovery indice for immobility time. N = 10 mice in the LPC group for female mice and N= 9-10 mice in the LPC group for male mice; N = 10 mice in the LPC + rTSMS group for female mice and N = 9-10 in the LPC + rTSMS group for male mice. The dotted line represents pre-lesion values. Statistical analyses were performed using Mann-Whitney test (*=P≤ 0.05; **=P≤ 0.01).

## DISCUSSION

To our knowledge, this is the first study to demonstrate the effects of rTSMS in a model of focal spinal cord demyelination and inflammation, both at the tissue level and in terms of functional recovery in mice. Indeed, only one recent study has investigated the effects of RMS in a model of cuprizone-induced demyelination ^29^. In this study, the authors mainly focused on the effects of this treatment on OLs and demonstrated that transcranial RMS can modulate either the number or the length of internodes, depending on whether RMS was applied during or after cuprizone exposure ^29^.

In our study, as a first step, it was important to characterize the kinetics of LPC-induced effects in the spinal cord (Figure S1). Although this model has already been described and defined as inducing a transient lesion ^19^, we needed to define time points at which differences between spontaneous recovery and rTSMS-induced recovery could be expected when testing a therapeutic intervention. Our results showed that LPC injection induced strong inflammation and demyelination at 8 and 15 days post-injection, with a marked resolution of these parameters by day 15 (Figure S1). Locomotor deficits, in turn, peaked at day 3 post-injection before markedly improving by day 7 (Figure S4). These findings allowed us to define the relevant time points for our study, namely 8 and 15 days post-injection, to investigate the effects of rTSMS in our first set of experiments on spinal tissue and from D2 to D14 for locomotor recovery.

In this first set of experiments, we investigated the effects of rTSMS when the treatment was initiated the day after LPC injection. In this context, we demonstrate that rTSMS reduces demyelination, not only by limiting the demyelinated area but also by decreasing the accumulation of myelin debris (Figure 1). We also show that rTSMS decreases the size of the inflammatory area within the spinal parenchyma (Figure 1). These results can be explained by recent findings in an SCI model in which the authors proved that rTSMS stimulates the phagocytosis of myelin debris by macrophages ^30^.

Furthermore, our results provide new insights into the effects of rTSMS on the fibroglial scar (Figure 2), a feature that is poorly described in inflammatory and/or demyelinating models. While a dense fibrotic core has been well documented in mouse models of traumatic SCI ^31,32^, the presence of such fibrosis has, to our knowledge, only recently been reported in the Experimental Autoimmune Encephalomyelitis (EAE) model and only one study described it in the focal LPC-induced demyelination model used here ^19,26,33^. Interestingly, the fibrosis observed following LPC injection is not identical to that found in traumatic models; after SCI, fibrosis exclusively fills the lesion core, whereas after LPC injection, it develops within the lesion area, but it is less dense and partially intermingles with astrocytes (Figure 2). As we previously demonstrated in SCI models, rTSMS also modulates this fibroglial scar by reducing the astrocyte-free area and decreasing fibrosis, inducing tissue repair after inflammatory SCI (Figure 2) ^14^. These results are consistent with those of Dorrier et *al*., who demonstrated that reducing fibrosis in EAE model promotes tissue repair ^33^. Our results also highlight that rTSMS induces tissue repair at an early stage, 8 days after LPC injection, but no longer exerts such effects 15 days after injection (Figures 1 and 2). It could be due to the fact that, by 15 days post-injection, LPC produces less pronounced alerration on the spinal tissue (Figure S1), and as a result, the effects of rTSMS become indistinguishable from those of spontaneous recovery. Indeed, the area values for the LPC + rTSMS group at 7 days are close to those found for the LPC group at 14 days (Figures 1 and 2).

Finally, in this first series of experiments, we show that rTSMS induces functional locomotor recovery in mice (Figure 3). In effect, the unrestrained locomotion of mice was assessed at different time points before and after LPC injection using the open-field test. It shows that rTSMS treatment increases the distance traveled by the animals and decreases their immobility time (Figure 3). Moreover, by calculating recovery indices, we demonstrated that the animals exhibited improved recovery indices for all parameters, namely the two mentioned above as well as average speed (Figure 3).

In contrast to SCI, where no sex-specific effects have been reported ^34^, demyelinating inflammatory lesions of the spinal cord are known to be more frequent in women and to display marked sex disparities ^27^. Moreover, the sex-specific effects in inflammatory models, such as those induced by LPC, have been barely explored, unlike in other models such as EAE ^27^. Therefore, we investigated whether the effects of rTSMS differed between male and female mice when the treatment was initiated the day after LPC injection (Figures S3 and S4). In this context, rTSMS induced similar effects in both male and female mice, for tissue repair and functional recovery (Figures S3 and S4, respectively).

Subsequently, given that similar outcomes in histological and locomotor analyses between male and female mice may mask sex-related differences in the underlying molecular mechanisms, we next performed RNASeq analyses while separating females and males (Figures 4 and 5, respectively). These analyses revealed that rTSMS modulates a broad range of biological processes, particularly inflammatory and immune pathways, but with differential regulation between females and males (Figures 4 and 5, respectively). To better understand these differences, we performed GSEA analyses based on the RNASeq data (Figure 6). These analyses confirmed that rTSMS differentially regulates inflammatory and immune processes in male and female mice. Specifically, rTSMS preferentially modulates innate and adaptive immune responses and TL–related pathways in females, whereas in males it more strongly affects innate immunity, neutrophil activity, and phagocytosis (Figure 6). These findings highlight the strong effects of rTSMS on immune response regulation and reveal a sex-dependent effect of this treatment. Notably, the impact of rTSMS on TL is particularly interesting. Indeed, a recent study has demonstrated that in both Alzheimer’s disease patients and the MPTP mouse model, rTSMS exerts its effects in a Treg-dependent manner ^35^. In particular, this study highlights that RMS treatment in Parkinsonian (MPTP model) mice increases dopaminergic neuron survival and improves motor parameters. However, these beneficial effects are lost following the administration of a Treg blocker ^35^.

These results encouraged us to further investigate the specific effects of RMS on immune cells *in vitro*, by performing RNASeq analyses on microglia and macrophage cultures either exposed or not to the pro-inflammatory cytokine IL-1 (Figures 7, 8, and S5–S7). Overall, these *in vitro* results show that RMS modulates the transcriptional profile of microglia in the absence of IL-1 activation, whereas in contrast, RMS modulates the transcriptional profile of macrophages when they are activated by IL-1 (Figures 7 and 8). Interestingly, while control microglia and macrophages display a strong response upon IL-1 exposure (Figures S5 and S6, respectively), RMS attenuates IL-1 effects in macrophages and fully blocks them in microglia (Figure S7). These findings are consistent with other, albeit limited, reports in the literature showing that *in vitro* RMS exerts modulatory effects on inflammatory cells, particularly when these cells are in a pro-inflammatory state ^13,36,37^. These results on the inflammatory and immune response are particularly interesting and important, because although RMS appears to exert effects on various cell types, including neurons, astrocytes, or OLs, numerous studies across a wide range of models highlight its modulatory impact on inflammatory processes *in vivo* ^38^. Indeed, in recent years, many studies have investigated the effects of RMS in diverse models such as stroke, Alzheimer’s disease, Parkinson’s disease, traumatic brain injury, SCI, and anxiety- or depression-like models. Most of these studies report beneficial effects of RMS that are closely correlated with the regulation or modulation of inflammatory processes ^39–49^. Finally, in our study, to better approximate a preclinical context, we investigated the effects of RMS when applied at the peak of motor symptoms, i.e., 3 days after LPC injection (Figure S4). In this paradigm, unlike what we previously demonstrated, rTSMS exerted different effects between male and female mice (Figures 9 and 10). Specifically, under these conditions, rTSMS reduced inflammation and demyelination processes and modulated the fibroglial scar in female mice, but had no effect in males (Figure 9). Moreover, it induced moderate functional recovery in females but showed no beneficial effects in males (Figure 10). These experimental conditions highlight a strong sex-dependent effect of rTSMS in our model when treatment begins at the peak of motor deficits. In the literature, few studies have investigated sex-specific effects of RMS ^50,51^. However, one study has shown that RMS exerts sex-dependent effects depending on the stimulation pattern used and also on the age of the animals when treatment is applied in a stab wound injury model ^51^. The discrepancy we observe here may be related to intrinsic variations in immune and inflammatory responses between males and females. Indeed, a recent study in humans showed that non-responder patients to RMS in conditions such as depression are those who notably exhibit the highest neutrophil-to-lymphocyte ratio in the blood ^52^. It has been shown that this ratio is higher in males than in females, both in humans and in rodents ^53,54^. In our model, we can hypothesize that by day three the immune response is already established, and according to our GSEA data it is characterized by an innate, neutrophil-dependent response in male mice (Figure 6B). This could explain, in this context, the lack of response observed in males when RMS is applied three days after LPC injection (Figures 9 and 10). In light of these results, it would be particularly interesting to further investigate the effects of RMS on the immune response, but also the impact of immune status on responsiveness to RMS. Indeed, in a clinical perspective, these data are crucial as they could allow prediction of patient responsiveness to the treatment and the design of differentiated therapeutic approaches depending on the patient’s immune status, for instance by combining RMS with another immunomodulatory therapy.

### Limitations and conclusion

As previously mentioned, our study is the first to investigate the effects of RMS in a focal demyelination model induced by LPC. However, although the results are very encouraging, they will need to be confirmed in other models of inflammatory SCI or in other models of spinal cord demyelination, given the kinetics of the LPC model. Indeed, although easy to implement, this model induces only a transient demyelination of the spinal cord, which spontaneously resolves after 4 weeks, as described in the literature and as shown in Figure S1 ^19^. It would therefore be necessary to test the effects of this treatment in other models of inflammatory lesions of the nervous system. In addition, the literature reports a wide variety of RMS protocols ^13^, and we can hypothesize that although our protocol proved effective in this model, other paradigms may yield better outcomes or modulate additional physiological parameters beyond those identified in our study. Finally, as previously mentioned, RMS appears to exert particularly important effects on the inflammatory and immune responses. Continuing investigations in this direction is essential to gain a deeper understanding of this treatment and to propose its use in a rational and clinically relevant manner. Future studies will also need to take into account sex-related differences, which remain poorly documented, particularly in the context of responses to neuromodulation-based therapies.

To conclude, our study demonstrates that rTSMS non-invasively reduces inflammation and demyelination in a focal demyelinating inflammatory model of the spinal cord in a sex-dependent manner. These findings provide new insights into the mechanisms of rTSMS and offer hope for its potential application in the treatment of clinical inflammatory diseases in humans.

## FUNDING SOURCES

This research was supported by Fondation des gueules cassées (AP 28-2023, AP 38-2024 and AP 46-2025).

## USE OF AI

ChatGpt was used to correct the spelling, grammar and syntax of the sentences.

## CONSENT FOR PUBLICATION

All authors have read the manuscript and indicated consent for publication

## COMPETING INTERESTS

The authors declare that they have no competing interests.

## AUTHOR CONTRIBUTIONS

F.S., Q.D. and N.G. conceptualized the project.

F.S. and N.G. designed the experiments.

F.S., I.Z., A.D., I.I., L.D., L.M., P.N., C.R. and J.D. performed the experiments.

F.S., A.D., L.D. and N.G. analyzed the results.

F.S. and N.G. wrote the article.

## ETHICS APPROVAL

All procedures that involved animal experiments were approved by the Ethics Committee of Paris Cité University France on September 1, 2023 for a five years period, with approval number of #44348-2023071210223131 v3. Title of the approved project was “ Study of the Effects of Repetitive Trans-Spinal Magnetic Stimulation in an Experimental Mouse Model of Transverse Myelitis”.

---

**Figure S1:**
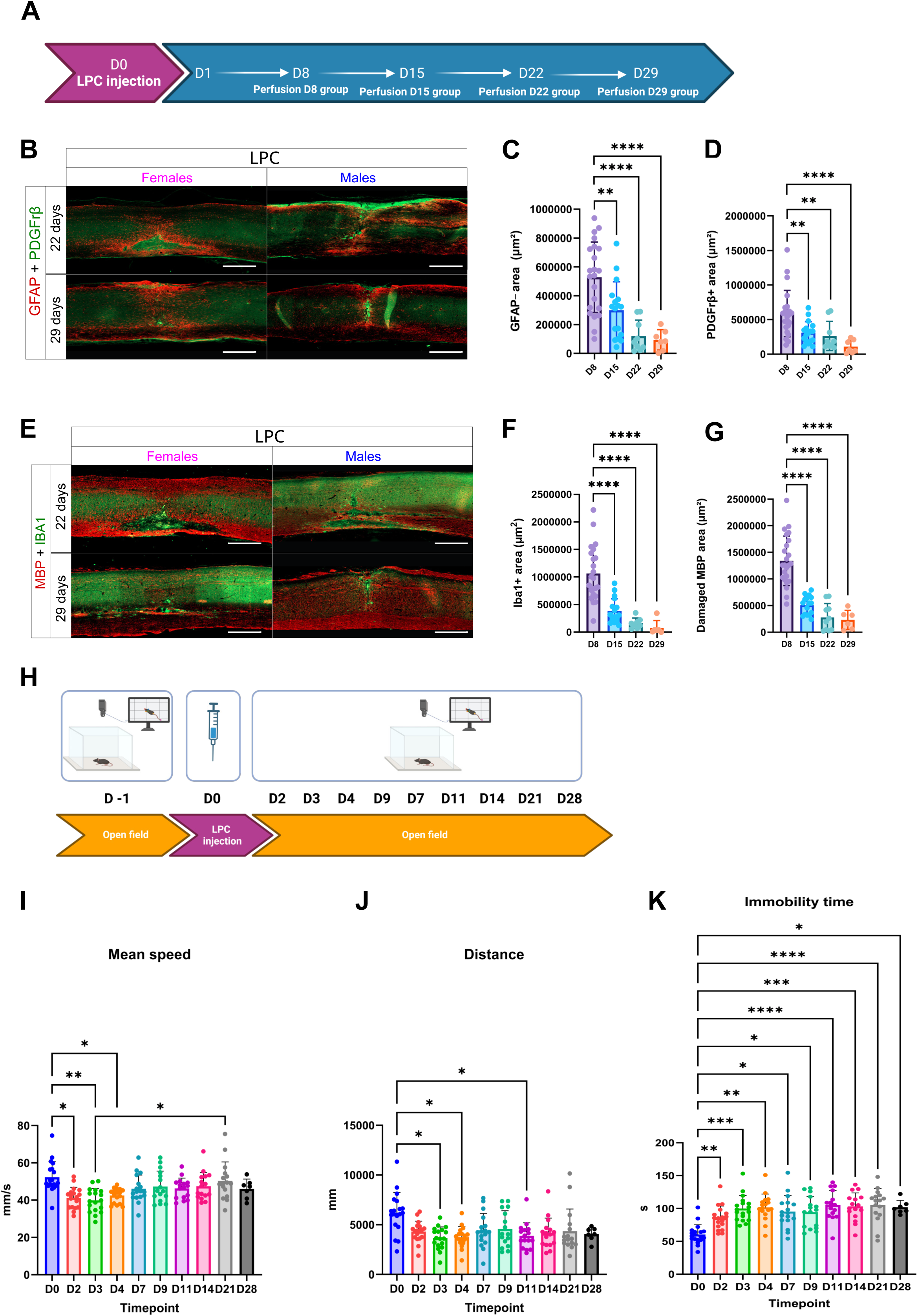
LPC injection induces transient inflammation, demyelination, fibrosis and astrocytic defect. **A.** Experimental design: on day 0, all mice received LPC injection and were perfused at D8, D15, D22 and D29 for histological analyses. **B-D.** Analysis of the effects of LPC on fibroglial scar formation 8, 15, 22 and 29 days after LPC injection. **B.** Representative pictures of sagittal spinal cord sections of LPC, males and females at 22 and 29 days after LPC injection. Sections were stained with anti-GFAP (in red) and anti-PDGFrβ (in green) antibodies. **C**. Quantification of GFAP-area. **D.** Quantification of PDGFrβ+ area. **E-G.** Analysis of the effects of LPC on inflammation and demyelination processes 8, 15, 22 and 29 days after LPC injection. **E.** Representative pictures of sagittal spinal cord sections of LPC males and females at 22 and 29 days after LPC injection. Sections were stained with anti-MBP (in red) and anti-Iba1 (in green) antibodies for Iba1 and damaged MBP area analyses. **F**. Quantification of Iba1+ area. **G.** Quantification of damaged MBP area (MBP defect + MBP debris areas). **H.** Experimental design: on day 0, all mice received LPC injection, then locomotor behaviors were recorded for all mice one day before surgery as well as on D2, D3, D4, D7, D9, D11, D14, D21 and D28. **I-K.** Analysis of the effects of LPC on functional recovery in a free-field locomotion - open field test analysis over time (from D0 to 28 days post-LPC injection). **I.** Quantification of mean speed. **J.** Quantification of distance traveled. **K.** Quantification of immobility time. N = 18 mice females + males in the LPC group. Statistical analyses were performed using Welch’s t-test (*=P≤ 0.05; **=P≤ 0.01; ***=P≤ 0.001 and ****=P≤ 0.0001).

**Figure S2:**
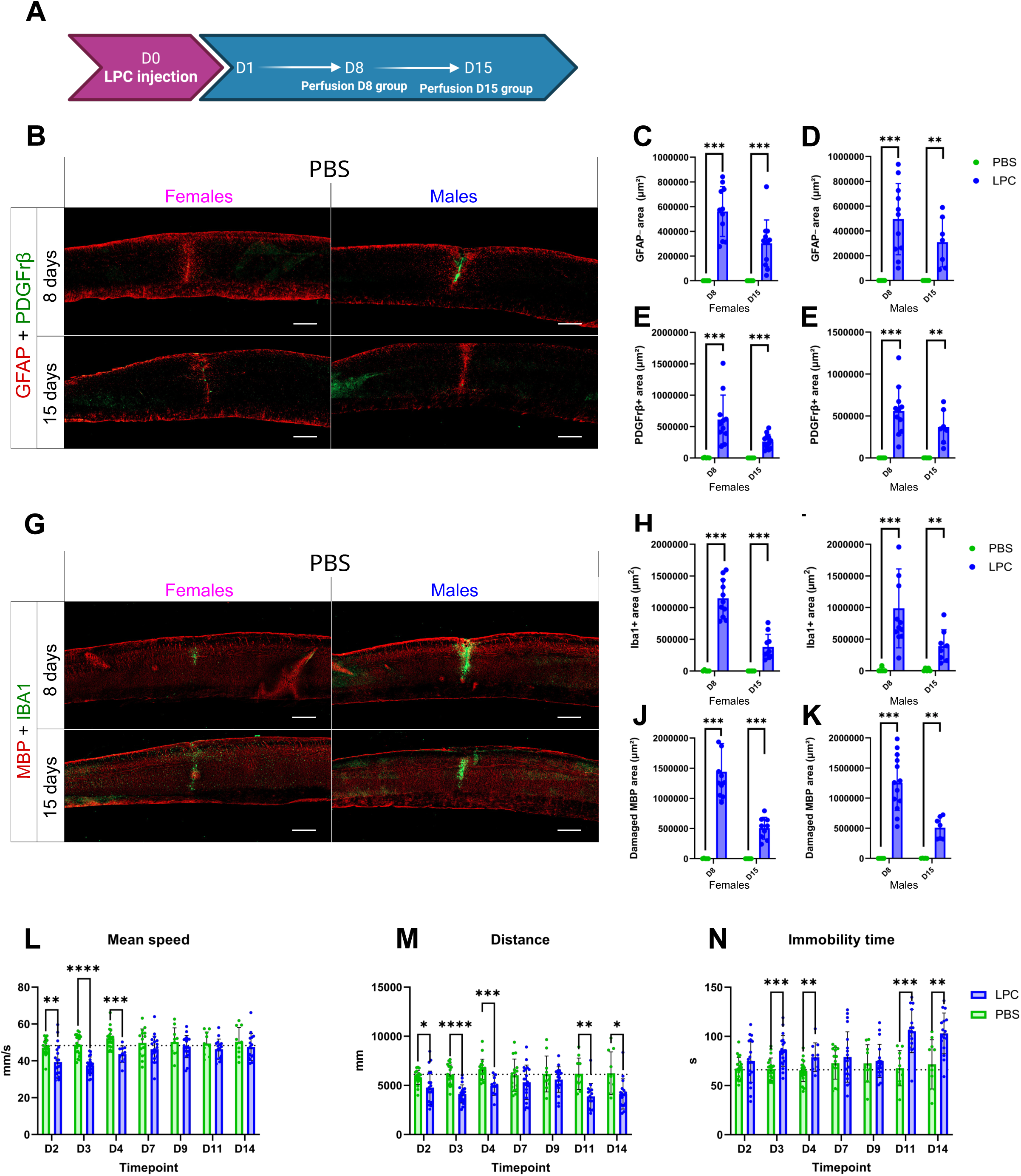
PBS injection does not impair locomotor function and does not induce fibrosis, astrocytic defect, demyelination and inflammation. **A.** Experimental design: on day 0, all mice received PBS or LPC injection and all mice were perfused at D8 and D15 for histological analyses. **B-F.** Analysis of the effects of PBS on fibroglial scar formation 8 and 15 days after PBS or LPC injection with PBS mice in green and LPC mice in blue. **B.** Representative pictures of sagittal spinal cord sections of PBS in male and female mice at 8 and 15 days after PBS injection. Sections were stained with anti-GFAP (in red) and anti-PDGFrβ (in green) antibodies. **C**. Quantification of GFAP-area in the female group. **D**. Quantification of GFAP negative area in the male group. **E.** Quantification of PDGFrβ+ area in the female group. **F.** Quantification of PDGFrβ+ area in the male group. **G-K.** Analysis of the effects of PBS on inflammation and demyelination processes 8 and 15 days after PBS or LPC injection with PBS mice in green and LPC mice in blue. **G.** Representative pictures of sagittal spinal cord sections of PBS in male and female mice at 8 and 15 days after PBS injection. Sections were stained with anti-MBP (in red) and anti-Iba1 (in green) antibodies for Iba1 and damaged MBP area analyses. **H**. Quantification of Iba1+ area in the female group. **I**. Quantification of Iba1+ area in the male group. **J.** Quantification of damaged MBP area (MBP defect + MBP debris areas) in the female group. **K.** Quantification of damaged MBP area (MBP defect + MBP debris areas) in the male group. N = 5 females and N = 5 males mice in the PBS group and N = 11 females and N = 11 males in the LPC group. Statistical analyses were performed using Mann-Whitney test (**=P≤ 0.01 and ***=P≤ 0.001). **L-N.** Analysis of the effects of PBS on functional recovery in a free-field locomotion - open field test analysis over time (from D0 to 14 days post-PBS or LPC injection) with PBS mice in green and LPC mice in blue. **L.** Quantification of mean speed. **M.** Quantification of distance traveled. **N.** Quantification of immobility time. The dotted line represents pre-lesion values. N = 20 mice in the PBS group at D2, D3, D4 and D7 ; N = 10 mice in the PBS group at D9, D11 and D14; N = 20 mice in the LPC group at D2, D3, D4 and D7 and N = 14 mice in the LPC group at D9, D11 and D14. Statistical analyses were performed using Welch’s t-test (*=P≤ 0.05; **=P≤ 0.01; ***=P≤ 0.001 and ****=P≤ 0.0001).

**Figure Supp 3:**
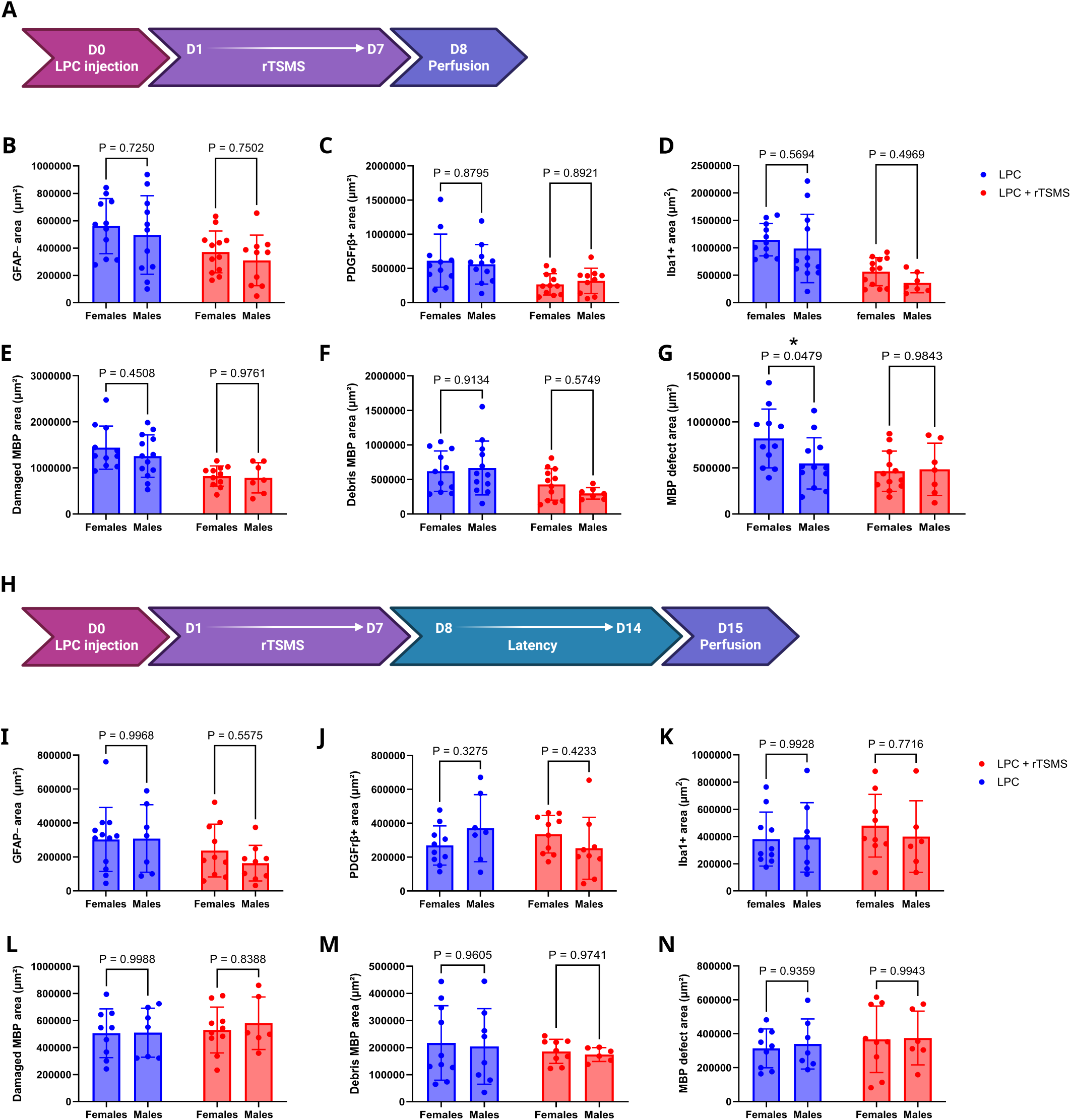
LPC injection and rTSMS treatment elicit similar tissue responses in male and female mice when applied at presymptomatic stage. **A.** Experimental design: on day 0, all mice received LPC injection, half of them underwent rTSMS treatment from D1 to D7 and all mice were perfused at D8 for histological analyses. **B-G.** Analysis of the sex-effect on fibroglial scar formation, inflammation and demyelination processes 8 days after LPC injection or rTSMS treatment in female and male mice with LPC group in blue and LPC + rTSMS group in red. **B**. Quantification of GFAP-area. **C.** Quantification of PDGFrB+ area. **D**. Quantification of Iba1+ area. **E.** Quantification of damaged MBP area (MBP defect + MBP debris areas). **F**. Quanfitication of MBP debris area. **G.** Quantification of MBP defect (MBP-area). **H.** Experimental design: on day 0, all mice received LPC injection, half of them underwent rTSMS treatment from D1 to D7 and D15 for histological analyses. **I-N.** Analysis of the sex-effect on fibroglial scar formation, inflammation and demyelination processes 15 days after LPC injection or rTSMS treatment in female and male mice with LPC group in blue and LPC + rTSMS group in red. **I**. Quantification of GFAP-area. **J.** Quantification of PDGFrB+ area. **K**. Quantification of Iba1+ area. **L.** Quantification of damaged MBP area (MBP defect + MBP debris areas). **M**. Quanfitication of MBP debris area. **N.** Quantification of MBP defect (MBP-area). N = 11 mice in LPC group for female mice and N= 7 mice in LPC group for male mice at 8 days ; N = 10 mice in LPC + rTSMS group for female mice and N = 9 in LPC + rTSMS group for male mice at 8 days. N = 10 mice in LPC group for female mice and N= 7 mice in LPC group for male mice at 15 days; N = 10 mice in LPC + rTSMS group for female mice and N = 9 in LPC + rTSMS group for male mice at 15 days. Statistical analyses were performed using Mann-Withney test.

**Figure S4:**
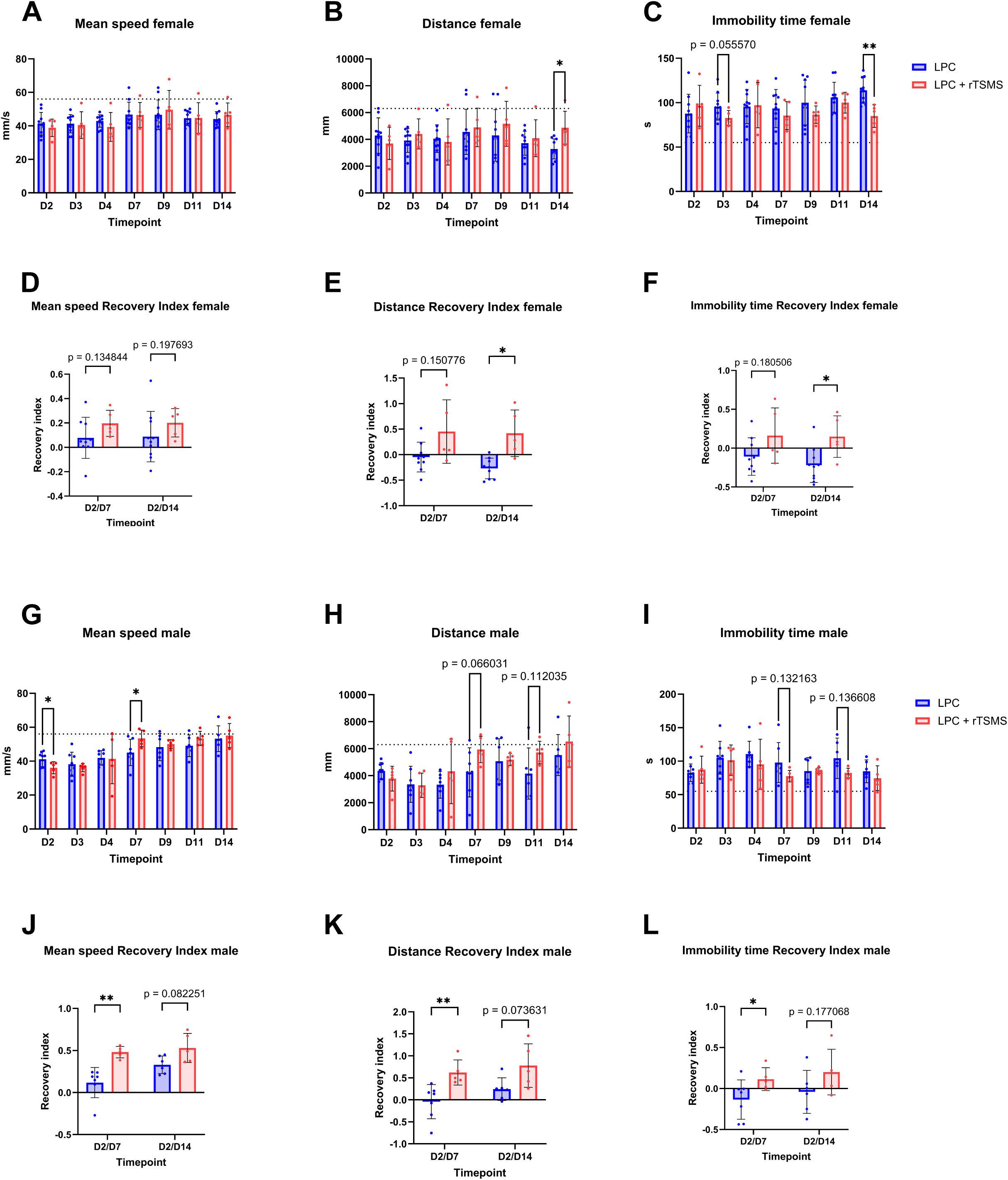
rTSMS treatment induces similar functional recovery in male and female mice. **A-C.** Analysis of the effects of rTSMS on functional recovery in a free-field locomotion - open field test analysis over time (from 2 to 14 days post-LPC injection) with LPC mice in blue and LPC + rTSMS mice in red in female mice**. A.** Quantification of mean speed. **B.** Quantification of distance traveled. **C.** Quantification of **i**mmobility time**. D-F.** Analysis of the effects of rTSMS on recovery indices with LPC mice in blue and LPC + rTSMS mice in red. Recovery indices were calculated for each animal by comparing the values measured at day 2 (D2) to those measured at day 7 (D7) and D14 (D14) post-lesion in female mice. **D.** Recovery indice for mean speed. **E.** Recovery indice for distance traveled. **F.** Recovery indice for immobility time. **G-I.** Analysis of the effects of rTSMS on functional recovery in a free-field locomotion - open field test analysis over time (from 2 to 14 days post-LPC injection) with LPC mice in blue and LPC + rTSMS mice in red in male mice**. G.** Quantification of mean speed. **H.** Quantification of distance traveled. **I.** Quantification of immobility time**. J-L.** Analysis of the effects of rTSMS on recovery indices with LPC mice in blue and LPC + rTSMS mice in red in male mice. Recovery indices were calculated for each animal by comparing the values measured at day 2 (D2) to those measured at day 7 (D7) and D14 (D14) post-lesion. **J.** Recovery indice for mean speed. **K.** Recovery indice for distance traveled. **L** Recovery indice for immobility time. N = 10 mice in the LPC group for female mice and N= 6 mice in the LPC group for male mice; N = 5 mice in the LPC + rTSMS group for female mice and N = 5 in the LPC + rTSMS group for male mice. The dotted line represents pre-lesion values. Statistical analyses were performed using Welch’s t-test (*=P≤ 0.05; **=P≤ 0.01).

**Figure S5:**
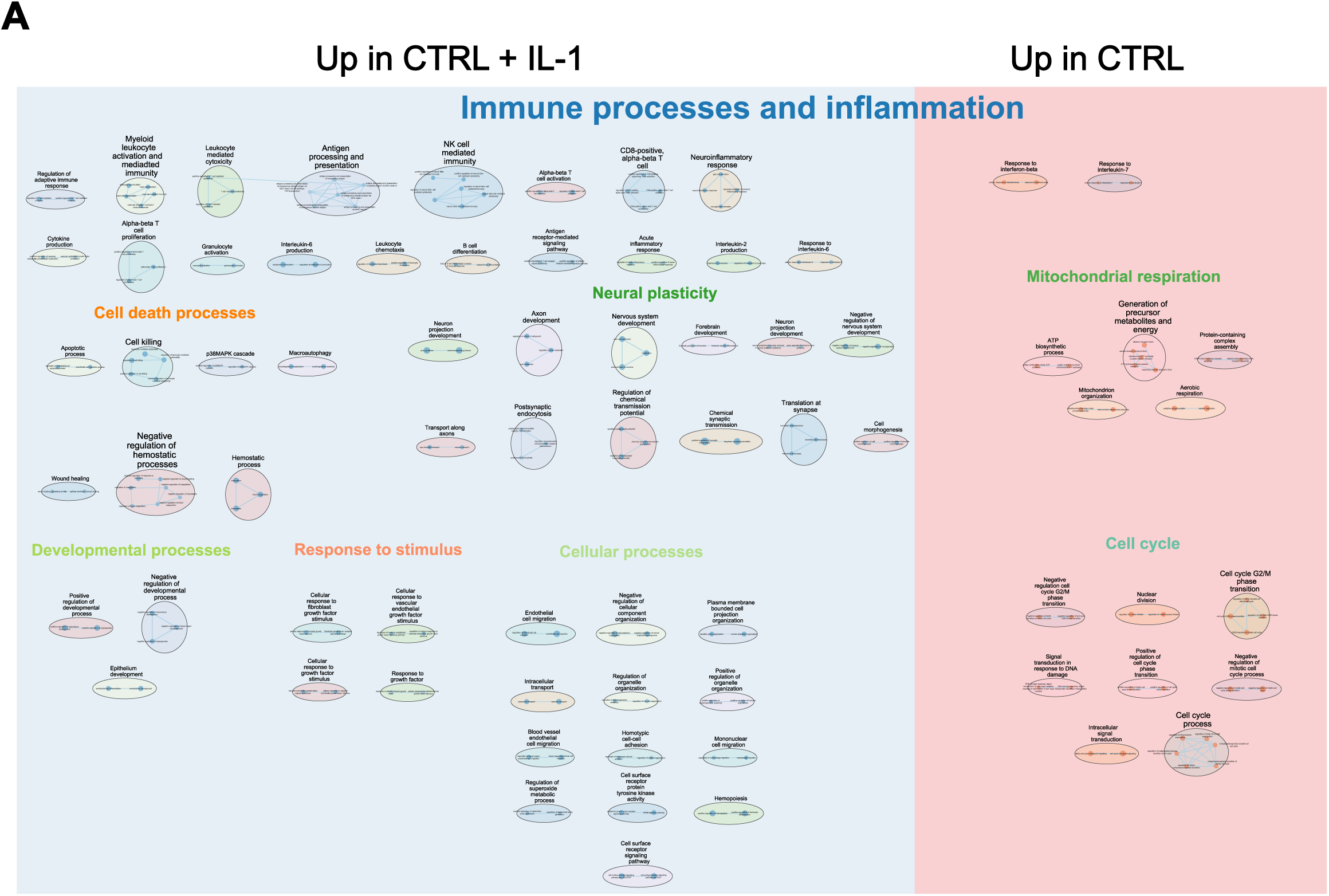
IL-1 modulate gene expression in BV2 cell cultures. **A.** Cytoscape representation of deregulated GO terms (“Biological Process” category). Each dot represents a deregulated GO-Term; their size and color are, respectively, proportional to the number of genes in the GO-term and the enrichment adjusted p-value (false discovery rate q value). Blue indicates GO terms upregulated in the CTRL + IL-1 group, while red indicates those upregulated in the CTRL group. GO-Terms are grouped into categories using the Autoannotate Cytoscape app (plugin). N = 5 independents samples in the CTRL group and N = 5 independents samples in the CTRL + IL-1 group.

**Figure S6:**
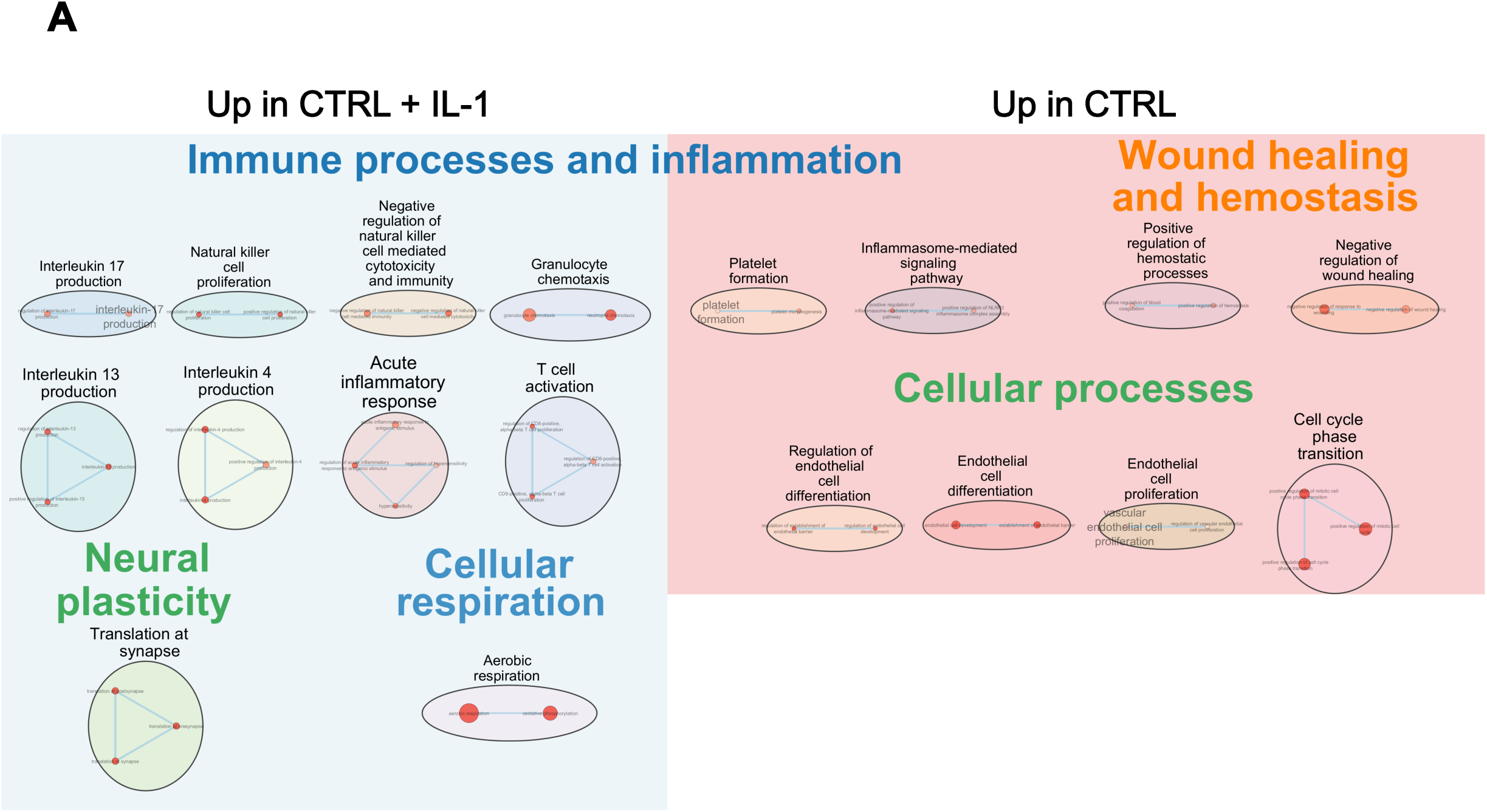
IL-1 modulate gene expression in RAW cell cultures. **A.** Cytoscape representation of deregulated GO terms (“Biological Process” category) s. Each dot represents a deregulated GO-Term; their size and color are, respectively, proportional to the number of genes in the GO-term and the enrichment adjusted p-value (false discovery rate q value). Blue indicates GO terms upregulated in the CTRL + IL-1 group, while red indicates those upregulated in the CTRL group. GO-Terms are grouped into categories using the Autoannotate Cytoscape app (plugin). N = 5 independents samples in the CTRL group and N = 5 independents samples in the CTRL + IL-1 group.

**Figure S7:**
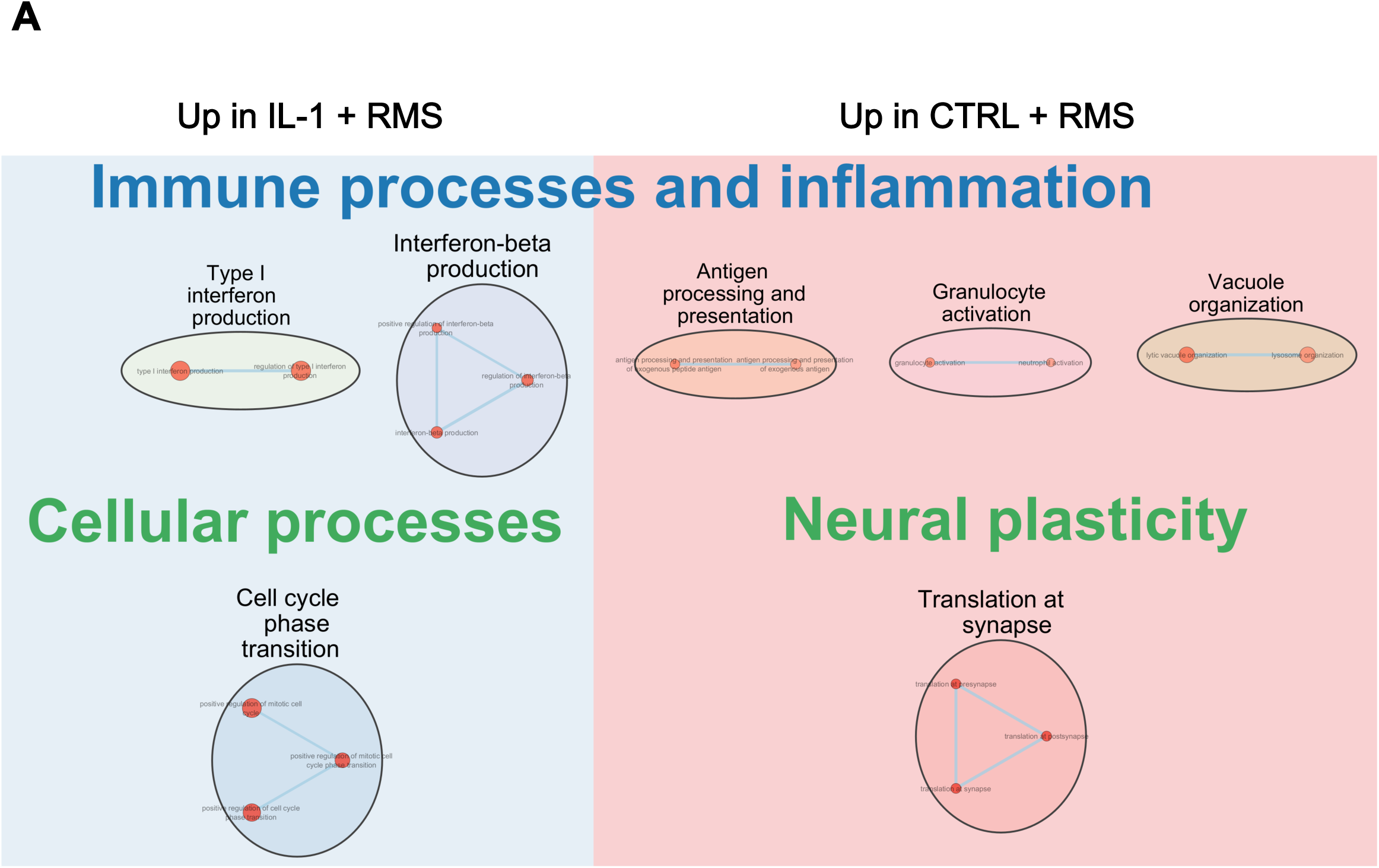
RMS attenuates IL-1 effects in microglia. **A.** Cytoscape representation of deregulated GO terms (“Biological Process” category). Each dot represents a deregulated GO-Term; their size and color are, respectively, proportional to the number of genes in the GO-term and the enrichment adjusted p-value (false discovery rate q value). Blue indicates GO terms upregulated in the IL-1 + RMS group, while red indicates those upregulated in the CTRL + RMS group. GO-Terms are grouped into categories using the Autoannotate Cytoscape app (plugin). N = 5 independents samples in the RMS group and N = 5 independents samples in the RMS + IL-1 group.

## Notes

### Competing Interest Statement

The authors have declared no competing interest.

